# Polyhydroxyalkanoate carbonosomes impact the activity and viability of growth-arrested *Pseudomonas aeruginosa* during nutrient fluctuations and tobramycin exposure

**DOI:** 10.64898/2026.10.04.756612

**Authors:** Findlay D Munro, Megan Bergkessel

**Affiliations:** Faculty of Life Sciences, University of Dundee, Dundee, UK

## Abstract

The opportunistic pathogen *Pseudomonas aeruginosa* can store carbon by polymerizing (R)-3-hydroxy fatty acids into polyhydroxyalkanoates (PHA) and forming large intracellular carbonosomes. These structures primarily function as stores of excess carbon that can be employed during nutrient imbalances, though they have been increasingly reported to have secondary functions in stress responses. A range of Proteobacteria can form carbonosomes, and they have been well characterised in strains used in bioplastic production and wastewater treatment. However, the occurrence, regulation and functions of carbonosomes in pathogenic bacteria remain poorly understood. Using flow cytometry and fluorescence microscopy following staining of carbonosomes with BODIPY 493/503, we characterised carbonosome abundance and dynamics in growth-arrested *P. aeruginosa* starved of nitrogen but provided with excess carbon. We found that carbonosomes substantially impact the energetic and biosynthetic output of growth-arrested *P. aeruginosa,* affecting protein biosynthesis, flagellar motility, and ATP/NADPH pools. The production and use of carbonosomes was found to confer a strong fitness advantage to growth-arrested *P. aeruginosa* during fluctuating nutrient conditions. Small carbonosomes were observed during the lag phase of growth in rich media and the ability to produce these structures was conserved across several different laboratory and clinical strains, although with different dynamics and maximal amounts. For example, the Liverpool epidemic strain LESB58 showed very little BODIPY staining while the closely related LESB65 strain showed high levels of staining. Crucially, we found that growth-arrested cells depolymerize carbonosomes upon tobramycin exposure and that carbonosome depolymerization significantly increases *P. aeruginosa* tobramycin sensitivity. Together, our observations suggest that carbonosomes significantly impact the physiology, metabolism, and antibiotic susceptibility of *P. aeruginosa*.

## Introduction

Laboratory studies into bacterial physiology have usually been conducted under conditions that permit exponential growth, where cells are assumed to exist in steady state, with concentrations of enzymes and metabolites remaining approximately constant [1, 2]. During the establishment of steady state exponential growth, regulatory networks adjust the allocation of proteome resources to maximize the growth rate [3–5]. However, in many environments, exponential growth cannot be sustained due to scarcity of one or more resources and the parameters to be optimized by regulatory networks are less clear. Cells may divert resources to stress responses that directly counter environmental insults [6, 7], rather than building biomass, or minimize energy expenditure and prolong survival by recycling internal macromolecules [8]. Additionally, they may store nutrients that are available in excess to prepare for future changes to nutrient availability [9–12]. The regulatory mechanisms impacting decisions to spend resources on one activity versus another during growth arrest or other non-steady state conditions are not well understood even in model organisms like *Escherichia coli* [13]. The physiological and evolutionary consequences of these decisions are even less explored, especially in opportunistic pathogens.

*Pseudomonas aeruginosa* is an opportunistic pathogen that causes difficult to treat infections in vulnerable patient populations, such as people with cystic fibrosis (pwCF), burn wounds, or indwelling medical devices [14]. However, *P. aeruginosa* is also ubiquitous in many natural environments such as water and soil [15]. In infections and other environments, nutrient depletion by the host organisms [16], competing microbes [17, 18], or clonemates in dense communities [19] can lead to low and unpredictable nutrient availability. Dynamic regulatory changes could confer survival advantages under these conditions despite an ongoing failure to achieve steady state exponential growth. To gain insight into these regulatory dynamics, we previously interrogated cellular physiology, mutant fitness, and new protein expression during a transition between nitrogen starvation and carbon starvation. We found that cells maintain flagellar motility and active synthesis of new proteins over days of nitrogen or carbon starvation, and that key global regulators such as the membrane associated protease FtsH, the global metabolic regulator PtsP, and the negative regulator of flagellar biosynthesis FleN, are important for fitness in both starvation conditions [20].

However, there were also obvious differences between cells under the two starvation conditions. Compared to carbon starvation, cells under nitrogen starvation were larger, contained fewer ribosomes, produced more new proteins, were more likely to continue swimming, and had higher optical density [20]. To further investigate this difference in optical density, which has been shown to be influenced by intracellular storage [21], we tested for the presence of previously described “carbonosomes”. These are aggregated polymers of polyhydroxyalkanoate (PHA) that can be stained and quantitatively measured using the fluorescent neutral lipid dye BODIPY 493/503 [22, 23]. Indeed, BODIPY staining revealed carbonosomes that occupied large fractions of the cytoplasmic area specifically during nitrogen starvation. We also found that carbonosome formation was affected by mutations in genes with starvation fitness impacts, as had been previously reported in *Pseudomonas putida* [24]. During nitrogen starvation, *ΔftsH* and *ΔptsP* mutant strains failed to produce carbonosomes, while a *flgE* transposon mutant (which shows a mild fitness advantage during the transition from nitrogen to carbon starvation) produced larger carbonosomes than the WT [20]. Together, these results suggest that carbonosomes might have broadly important consequences for cellular physiology under conditions outside of steady state growth, where carbon availability might fluctuate.

The pathways leading to carbonosome production have been studied in detail in non-pathogenic bacteria such as *P. putida* [25] and *Cupriavidus necator* [26]. In these species, and in the marine Proteobacteria *Phaeobacter inhibens* and *Vibrio splendidus,* carbonosomes have been shown to contribute to fitness during nutrient fluctuations [27–30]. However, previous work has often focused on maximizing yields of the specific PHB (polyhydroxybutyrate) polymer for use in bioplastic production [31, 32], rather than on the natural physiological roles of carbonosomes. In *P. aeruginosa* UCBPP-PA14, a 7-gene locus encodes structural, enzymatic and regulatory proteins that participate in carbonosome formation (Figure 1A). Two PHA polymerases, encoded by *phaC1* and *phaC2*, polymerize medium chain length (R)-3-hydroxy acyl-CoA, derived from fatty acid β-oxidation or *de novo* synthesis pathways, into PHA [25] (Figure 1B). Data from *P. putida* and *P. aeruginosa* suggest that *phaC1* is co-transcribed with the gene encoding the PHA depolymerase, *phaZ,* and regulated by housekeeping, stationary phase and nitrogen-responsive sigma factors in conjunction with the *phaD* transcription factor encoded immediately downstream [33–35]. The phasin genes *phaF* and *phaI* encode self-oligomerising proteins which form a proteinaceous shell surrounding the carbonosome [36], although PhaF has recently been described to have an additional role in translational regulation of many genes [37]. The function of *PA14_66890* is not known but it is associated with carbonosome genes in multiple pseudomonads [38].

**Figure 1.**
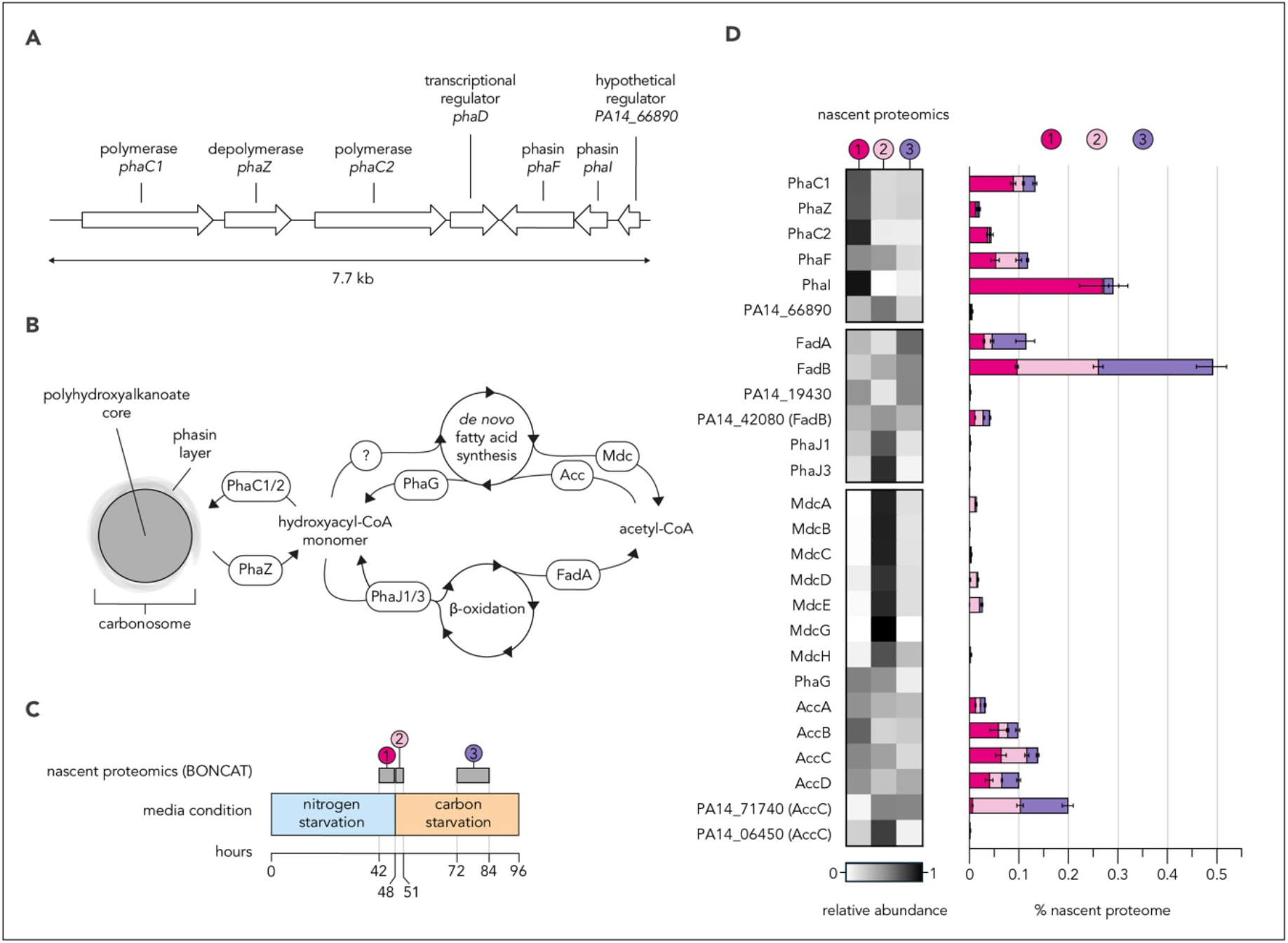
Trends in expression of carbonosome-associated proteins during nutritional fluctuation of *P. aeruginosa*. Diagram **(A)** shows architecture of the locus encoding enzymatic, structural and regulatory carbonosome-associated proteins in *P. aeruginosa* UCBPP-PA14. Arrows represent genes as annotated by pseudomonas.com [39], with length and directionality corresponding to gene length and transcription directionality. Each gene is labelled with common name or locus tag and functionality of its protein product. Diagram **(B)** shows the metabolic circuitry underlying the bidirectional conversion of acetyl-CoA to polyhydroxyalkanoate in *P. aeruginosa*. Where known, enzymes responsible for reactions are annotated. Schematic **(C)** shows the timeframe, starvation conditions and BONCAT labelling periods previously used to quantify carbonosome dynamics and inspect nascent proteome composition across a rapid shift from nitrogen-starvation to carbon-starvation minimal media [20]. Paired heat map (left) and bar chart (right) **(D)** show the abundance and dynamics of relevant nascent proteins identified by LC-MS/MS from the BONCAT labelling periods shown in diagram A [20]. Heat maps show the relative abundance of each nascent protein at each labelling period compared to the combined abundance across all three periods. Bar graphs show abundances as a percentage of the total nascent proteome at each labelling period. n=4 biological replicates for each labelling period.

In our previous investigation of the transition between nitrogen and carbon starvation, we used BONCAT (<u>B</u>io-<u>O</u>rthogonal <u>N</u>on-<u>C</u>anonical <u>A</u>mino Acid <u>T</u>agging) to show that a burst of new protein synthesis occurred immediately following a shift from nitrogen to carbon starvation, even though the average rate of protein synthesis during ongoing carbon starvation was very low [20]. The BONCAT method makes use of an azide-containing methionine analog (azidohomoalanine, AHA) that is incorporated into newly synthesized proteins during a chosen labelling period [40]. An azide-alkyne cycloaddition can then conjugate a fluorophore to AHA-containing proteins in fixed single cells, producing a signal proportional to the amount of new protein made during the labelling period [41]. We also used BONCAT to enrich and identify proteins that were newly synthesised before, during, and after the transition from nitrogen to carbon starvation by conjugating newly synthesized proteins from cell lysate to agarose beads and analyzing them by tryptic digest and LC-MS/MS [42–44] (Figure 1C).

Interestingly, we observed that several proteins predicted to be connected to carbonosome dynamics showed large changes in new synthesis during this transition (Figure 1D). The synthases PhaC1/C2, depolymerase PhaZ, transacylase PhaG, and phasin PhaI were most strongly expressed during nitrogen starvation conditions, and new synthesis decreased after the transition to carbon starvation, with PhaI showing one of the largest fold decreases of all nascent proteins [20]. All these proteins except PhaZ would be predicted to contribute to carbonosome production, but *phaZ* is co-transcribed with *phaC1* in *P. putida* [33]. The four subunits of the acetyl-CoA carboxylase (AccA, B, C, and D), which converts acetyl-CoA to malonyl-CoA, the direct substrate for *de novo* fatty acid synthesis, were also downregulated during the transition. In contrast, PhaJ1, PhaJ3, and the subunits of the malonyl decarboxylase complex (Mdc) were all sharply upregulated during the transition to carbon starvation, showing some of the largest fold increases of any proteins in the nascent proteome even if their absolute abundances were not very high (Figure 1D) [20]. PhaJ1 and PhaJ3 are enoyl-CoA hydratases that reversibly catalyze the conversion between (R)-3-hydroxyacyl-CoA and (R) enoyl-CoA, a substrate for β-oxidation of fatty acids which yields acetyl-CoA [45]. The Mdc complex converts malonyl-CoA into CO_2_ and acetyl-CoA and thus carries out the reverse reaction to the acetyl-CoA carboxylase complex[46]. Acetyl-CoA can be redistributed broadly throughout cellular metabolic pathways via the TCA cycle. Together, these observations suggested that changes to carbonosome dynamics are a meaningful component of the proteome remodelling that occurs upon a shift from nitrogen to carbon starvation.

In this work, we have explored the physiological roles of carbonosomes in *P. aeruginosa*, including their potential relevance in infection contexts. Using the neutral lipid dye BODIPY 493/503, we have tracked carbonosome accumulation and depolymerization during dynamic growth-arrested conditions. We generated mutants that lack the polymerases *phaC1* and *phaC2* and/or the depolymerase *phaZ*, yielding strains that completely lack the ability to produce carbonosomes, or lack the ability to depolymerize them. As previously reported for *C. necator* [27], we observed that these strains have fitness defects under conditions of fluctuating nutrient availability. We investigated the physiological impacts in more detail, showing that the inability to make or use carbonosomes resulted in changes to new protein synthesis (measured by BONCAT), ATP/NADPH levels (measured using a luciferase reporter), and flagellar motility (measured by tracking of individual cells), during a transition between nitrogen and carbon starvation.

Finally, we investigated whether the physiological and fitness impacts of carbonosomes might be relevant in infection contexts. Strikingly, we found that carbonosome depolymerization was triggered by treatment with the aminoglycoside antibiotic tobramycin, which is frequently used to treat chronic lung infections in pwCF [47]. This depolymerization correlated with increased ATP/NADPH pools and increased sensitivity to tobramycin. As with many virulence traits that can confer both advantages and disadvantages in heterogeneous infection contexts [48–50], we found that carbonosome production varied widely across a small selection of *P. aeruginosa* isolates. Together, this work suggests that carbonosomes can play important roles in *P. aeruginosa* under conditions that prevent steady state growth, with implications for both infection contexts and environmental lifestyles.

## Results

### *Pseudomonas aeruginosa* produces large carbonosomes under nitrogen starvation conditions

To study carbonosome formation over prolonged starvation time courses, we used conditions previously shown to preserve viability during at least a week of nitrogen starvation [20]. We resuspended cells from LB overnight cultures at low density (OD_500nm_ = 0.2) in a MOPS-based minimal medium with 45 mM sodium succinate as a carbon source but no nitrogen source. We periodically sampled and fixed aliquots of these nitrogen-starved cultures and stained them with BODIPY 493/503. Per-cell fluorescence of BODIPY-stained samples was measured by flow cytometry and fluorescence microscopy to assess carbonosome content, variation and positioning within nitrogen-starved populations as time progressed (Figure 2A).

**Figure 2.**
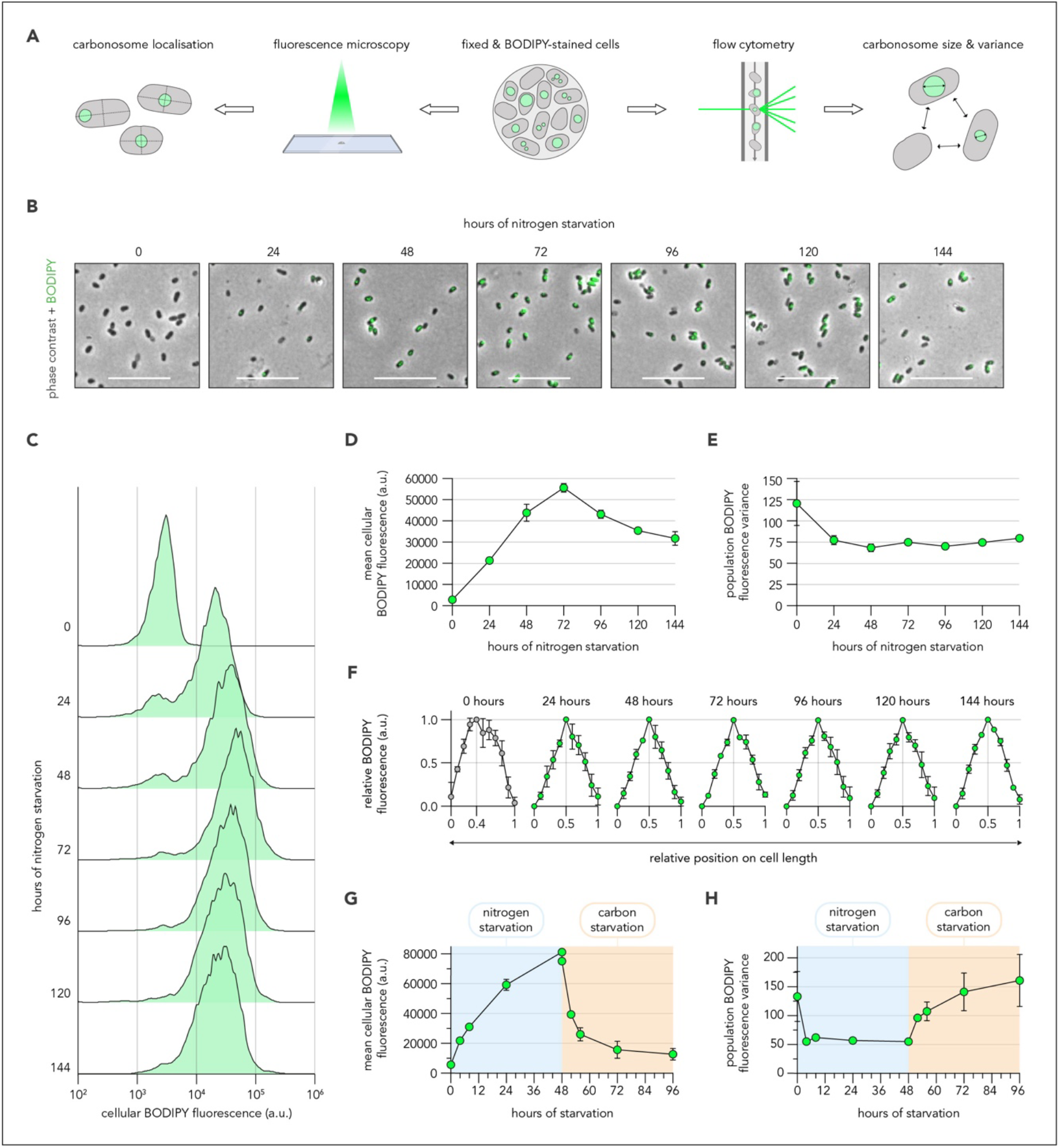
*P. aeruginosa* accumulates and depolymerizes carbonosomes under growth-limiting conditions. Schematic **(A)** shows experimental strategy for characterizing carbonosomes. Composite phase contrast and epifluorescence images **(B)** show fixed, BODIPY-stained cells from cultures incubated in nitrogen-starvation minimal media. Scale bar represents 10 µm. Histograms **(C)** show the flow cytometry measurements of per-cell BODIPY intensity for populations incubated in nitrogen starvation minimal media and sampled longitudinally. Line graphs show the mean per-cell fluorescence **(D)** and variance **(E)** extracted from cytometry-derived BODIPY-fluorescence histograms as shown in C. Line graphs **(F)** trace carbonosome positioning, as determined by fluorescence microscopy, along the axes of cells incubated in nitrogen starvation minimal media. Fluorescence values at each timepoint were normalized to the highest average fluorescence recorded along the cell axes, with this maximum value being denoted on respective x-axes. Grey points indicate that carbonosomes were not detected. Line graphs show the mean cellular BODIPY fluorescence **(G)** and variance **(H)** extracted from flow cytometry data of populations during a transition from nitrogen to carbon starvation. Panels D-H show means and standard deviations from n=3 biological replicates.

Using fluorescence microscopy, we saw that nitrogen starved cells possessed large, round regions of bright BODIPY fluorescence, which we interpret to be carbonosomes (Figure 2B, Supplemental figure 1). Flow cytometry was used to quantify BODIPY signal from thousands of cells, allowing assessment of the population distribution of BODIPY signal (Figure 2C) from which mean and variance values (Figure 2D, E), could be extracted. The distributions were largely unimodal. A small sub-population of low-fluorescence cells was notable at earlier timepoints, although it did not substantially affect the population variance (Figure 2C, E). The average BODIPY fluorescence of nitrogen starved populations dramatically increased over the first 72 hours of starvation and then modestly decreased (Figure 2C, D), matching the quantitative assessment from microscopy (Figure 2B, Supplemental Figure 1). The decrease in signal after 72 hours could be due to increasing depolymerase activity or decreasing polymerase activity, possibly influenced by ongoing depletion of succinate from the medium. Indeed, we observed an earlier depolymerization phase when the nitrogen starvation period was started at higher culture densities or with lower sodium succinate concentrations (Supplemental Figure 2/3). Fluorescence microscopy was also used to assess the average positioning of carbonosomes within starved cells by pinpointing the BODIPY fluorescence maxima along each cell length at each timepoint. On average, nitrogen-starved cells exhibited a single carbonosome which was centrally positioned throughout the time course (Figure 2F).

Having defined carbonosome dynamics in *P. aeruginosa* under protracted nitrogen-starvation conditions, we next asked how these structures would be impacted by sudden transition to carbon starvation. We exposed cells to nitrogen starvation conditions for 48 hours before shifting them to carbon starvation medium. (Figure 2G). Our previous investigations of this transition between nitrogen and carbon starvation showed that cells failed to detectably grow, suggesting that either nitrogen or carbon were limiting throughout [20]. Immediately following the transition, the mean cellular BODIPY fluorescence of populations decreased sharply (Figure 2G). More than half of the fluorescence signal was lost in the first 4 hours, and fluorescence dropped to near background levels 48 hours after the transition. Interestingly, population fluorescence variance increased following the transition, implying heterogeneity in the rate of carbonosome depolymerization (Figure 2H).

These findings suggest that *P. aeruginosa* can adjust carbonosome content in response to both gradual and rapid shifts in extracellular carbon availability.

### Carbonosome production and depolymerization enhance *P. aeruginosa* fitness during nutrient fluctuations

To test physiological impacts of both storing and releasing carbon from carbonosomes, we produced unmarked deletion strains which lacked the depolymerase PhaZ (Δ*phaZ*) or lacked both polymerases and the depolymerase (Δ*phaC1/Z/C2*) (Figure 3A). We first measured carbonosome production in these strains using both flow cytometry and microscopy to confirm expected phenotypes: we were unable to detect carbonosomes in the Δ*phaC1/Z/C2* mutant strain while Δ*phaZ* strain continuously increased in cellular BODIPY signal over the entire nitrogen starvation time course (Figure 3B/C, Supplemental Figure 3). Interestingly, the fluorescence histograms of Δ*phaZ* populations were narrow and lacked the small sub-population of carbonosome-lacking cells seen in the early hours of WT accumulation. Furthermore, when transitioned to carbon starvation conditions, the Δ*phaZ* strain failed to depolymerize carbonosomes (Figure 3D). These data suggest that activity of the depolymerase PhaZ contributes to the population heterogeneity observed in carbonosome signal in the WT during nitrogen starvation, in addition to being required for depolymerizing carbonosomes during carbon starvation.

**Figure 3.**
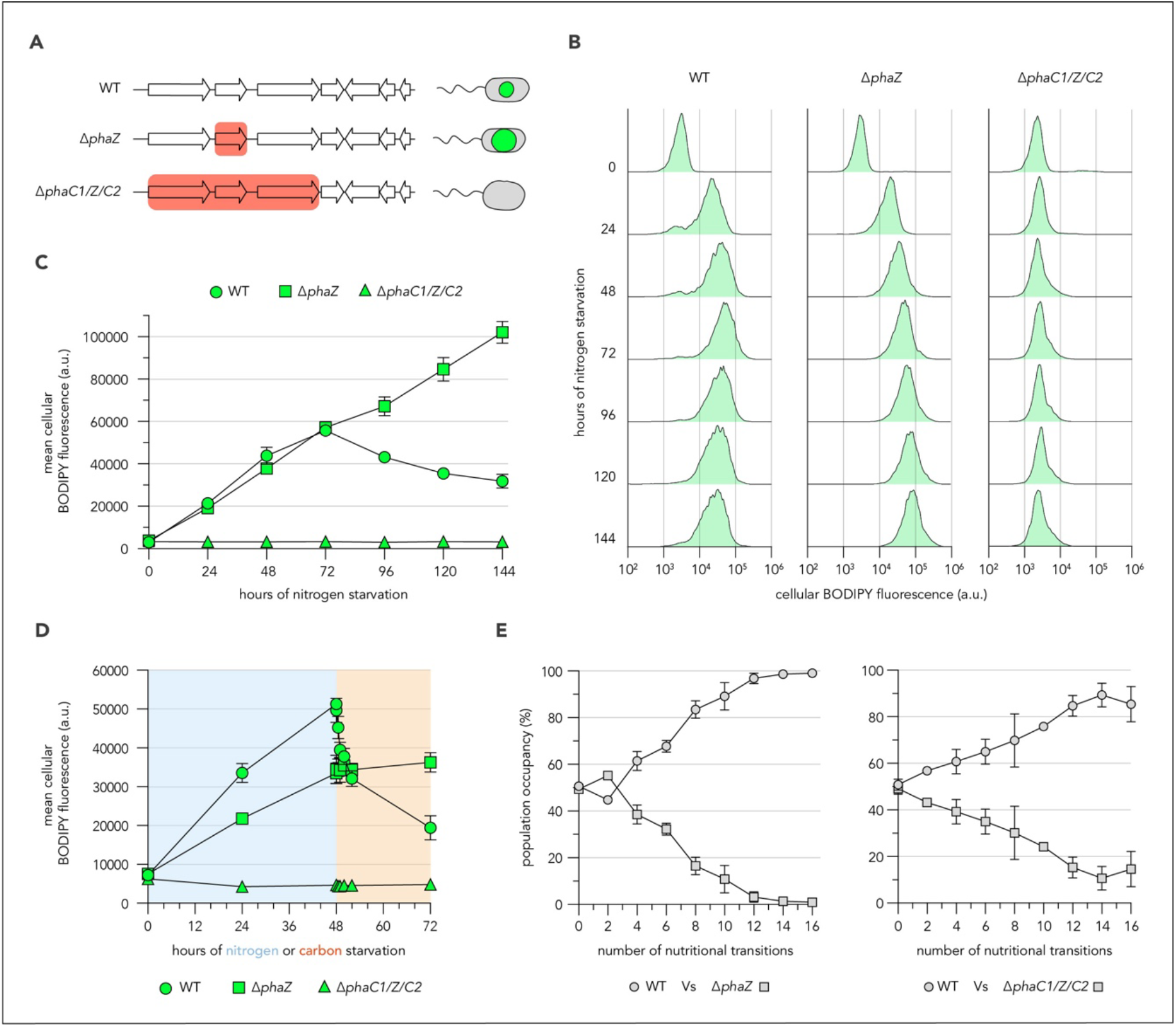
Carbonosome dynamics enhance *P. aeruginosa* fitness during nutrient fluctuations. Schematic **(A)** shows genotypes and expected phenotypes of WT and mutant *P. aeruginosa* strains used in subsequent investigations. White arrows represent individual genes of the carbonosome-associated locus annotated in 1A, with shaded red blocks denoting regions which are absent from unmarked deletion strains. Histograms **(B)** show the BODIPY fluorescence intensity, retrieved by flow cytometry, of WT, Δ*phaZ* and Δ*phaC1/Z/C2* populations incubated in nitrogen starvation minimal media and sampled longitudinally. Line graph **(C)** shows the mean cellular fluorescence extracted from BODIPY-fluorescence histograms of WT, Δ*phaZ* and Δ*phaC1/Z/C2* cells shown in B. Line graph **(D)** shows the mean cellular fluorescence extracted from cytometry-derived BODIPY-fluorescence histograms of WT, Δ*phaZ* and Δ*phaC1/Z/C2* populations incubated in nitrogen starvation minimal media for 48 hours before being transitioned to carbon starvation minimal media. Line graphs **(E)** show the average percentage of the mixed population in a competitive transition assay occupied by either strain across successive transitions between nitrogen-starvation and carbon-starvation minimal media. Panels C-E show means and standard deviations from n=3 biological replicates.

Despite their dramatic carbonosome phenotypes, both mutant strains grew in LB medium and survived nitrogen starvation similarly to the WT strain (Supplemental Figure 4). The optical densities (OD_500nm_) of the nitrogen-starved mutant cultures differed, with Δ*phaZ* having higher OD_500nm_ and Δ*phaC1/Z/C2* lower OD_500nm_ than WT. These optical density differences were not due to secretions into the supernatant, but instead increased opacity of the cells (Supplemental Figure 4), which has previously been observed in other carbonosome-producing microorganisms [21].

To test whether carbonosomes could confer fitness benefits to *P. aeruginosa*in fluctuating conditions, we mixed WT and mutant cells labelled with superfolder GFP or mScarlet-i (constitutively expressed from the chromosomal *attB* site) at a 50:50 ratio and repeatedly transitioned them between nitrogen and carbon starvation conditions. Previously, we did not observe a strong fitness effect of transposon insertion mutations in any of the carbonosome-associated genes in a Tn-Seq experiment that investigated a single transition from nitrogen to carbon starvation [20], but we wondered whether the benefits of carbonosomes might accrue over multiple transitions between these states. Using fluorescence microscopy, we determined the proportion of cells of each strain in the mixture following every second transition. We incubated mixtures for 48 hours after each transition to allow for carbonosomes to accumulate and depolymerise in WT cells, and we found that fluorophore choice did not substantially impact fitness measurements (Supplemental Figure 4). Successive transitions resulted in the continual loss of Δ*phaZ* and Δ*phaC1/Z/C2* mutant cells from mixed populations (Figure 3E). Interestingly, Δ*phaZ* cells were more quickly lost from mixed populations compared to Δ*phaC1/Z/C2* cells, suggesting that during fluctuating conditions the irreversible investment of material in carbonosomes more severely impacted viability or fitness than the lack of investment in carbonosomes. These data show that the ability to store and use carbonosomes during transitions between nitrogen and carbon starvation has a measurable fitness benefit.

### Carbonosomes impact the translational capacity, ATP/NADPH levels and flagellar motility of non-growing *P. aeruginosa*

We next probed mechanisms by which carbonosome utilization could support viability and fitness of growth-arrested *P. aeruginosa* during nutrient fluctuations. We and others previously showed that starved cells maintain low levels of ongoing new protein synthesis and flagellar motility [20, 41, 51], which requires continual replenishment of cellular ATP pools and reducing power. We hypothesized that the maintenance of these activities could have fitness benefits and that carbon liberated from carbonosomes could contribute to the proton motive force (which powers motility), amino acid pools, and ATP/NAD(P)H pools, directly and via acetyl-CoA incorporated into the TCA cycle. To test whether carbonosome production and utilization impacted these activities in growth-arrested *P. aeruginosa* we employed our Δ*phaZ* and Δ*phaC1/Z/C2* mutant strains in assays to measure new protein synthesis via BONCAT; ATP/NADPH pools via a luciferase reporter; and flagellar motility via single-cell tracking of timelapse microscopy.

We used sequential two-hour pulses of BONCAT labelling, followed by flow cytometry, to estimate per-cell new protein synthesis before, during, and after cultures were transitioned from nitrogen to carbon starvation conditions (Figure 4A). As previously observed [20], WT populations exhibited a burst of new protein synthesis immediately following the transition to the carbon starvation condition (Figure 4B), which aligned temporally with the carbonosome depolymerization also elicited by the transition (Figure 2G). Neither Δ*phaZ* nor Δ*phaC1/Z/C2* populations exhibited a burst of translation upon the transition to the carbon starvation condition. Instead, the average BONCAT signal of each mutant population rapidly decreased and remained low throughout the timecourse. These data suggest that robust carbonosome dynamics are required to support a burst of new protein synthesis in growth-arrested *P. aeruginosa* transitioning from nitrogen to carbon starvation conditions. The disruption of carbonosome dynamics in Δ*phaZ* and Δ*phaC1/Z/C2* cells could therefore hinder effective proteome remodelling in response to nutrient fluctuations.

**Figure 4.**
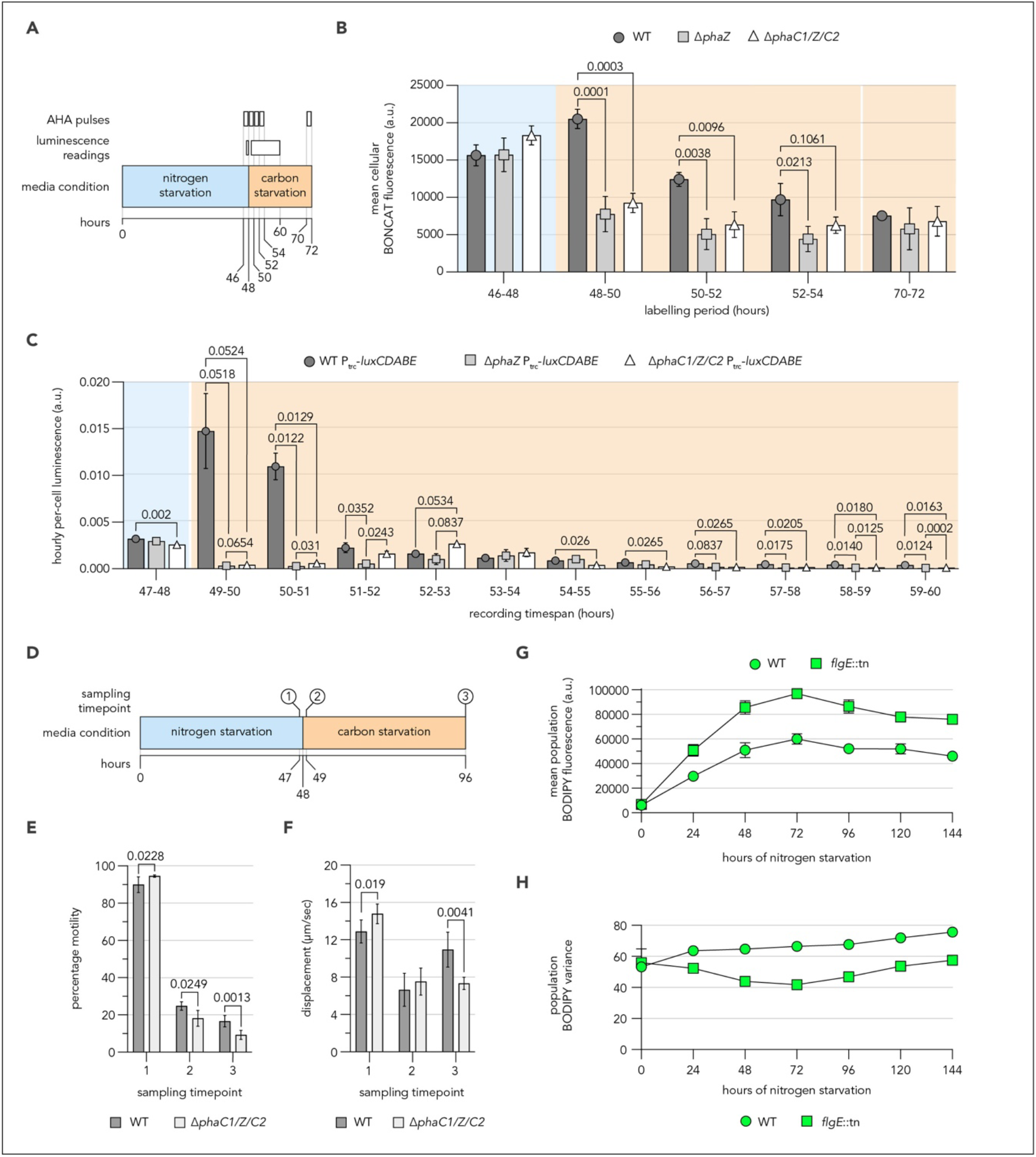
Carbonosomes impact the translational capacity, ATP levels and flagellar motility of non-growing *P. aeruginosa*. Schematic **(A)** shows the timeframe of BONCAT labelling pulses and luminescence readings during a transition from nitrogen-starvation to carbon-starvation minimal media. Bar graph **(B)** shows the average per-cell fluorescence from BONCAT labelling of cultures over the time course shown in A. Bar graph **(C)** shows the hourly summed luminescence signal (measured every 6 minutes) from cultures of P_trc_-*luxCDABE* strains during the time-course shown in A, normalised to CFU counts measured once in each medium. Schematic **(D)** shows the time course from which samples of culture were taken for time-lapse microscopy. Bar graphs show the percentage motility **(E)** and average speed of motile cells **(F)** from the time course shown in D. Line graphs show the mean per-cell fluorescence **(G)** and variance **(H)** of per-cell BODIPY signal from WT and *flgE*::tn cultures exposed to nitrogen starvation conditions. Means and standard deviations from n=3 biological replicates are shown. One-way ANOVAs with post-hoc Dunnett’s T3 multiple comparison or Welch’s unpaired T-tests were used to assess statistical significance for comparisons among 3 or 2 strains, with only values close to or passing significance thresholds (p < 0.05) shown.

In order to estimate changes in ATP/NADPH pools longitudinally over our transition time course (Figure 4A), we generated reporter strains in the WT and each mutant background in which the *luxCDABE* operon from *Photorhabdus luminescens* was constitutively expressed from the *attB* chromosomal location [52]. Previously, it has been shown that luminescence signal produced by these enzymes can be used to track metabolic fluxes in real time because cellular ATP/NADPH levels are rate limiting for regeneration of the fatty aldehyde substrate of the LuxAB luciferase by the LuxCE fatty acid reductase [53, 54]. The LuxCDABE-expressing strains were very similar to their parental strains in their growth in LB media, survival during nitrogen starvation, and ability to produce carbonosomes (Supplemental Figure 5). Summed luminescence values measured every six minutes during each hour before or after the transition between nitrogen and carbon starvation were calculated and normalized to colony forming unit (CFU) counts taken before or after the transition (Supplemental Figure 5).

The cellular luminescence signal of WT and carbonosome mutant *P. aeruginosa* were roughly comparable, (though significantly different due to low variance), under nitrogen starvation conditions (Figure 4C). However, following the transition to carbon starvation, the WT signal dramatically increased, rising clearly and significantly above the signal of the Δ*phaZ* and Δ*phaC1/Z/C2* mutants. Although both mutants later showed a slight increase in signal, this delayed response was much smaller than that observed in the WT cells. Crucially, all three strains behaved similarly in control transitions from nitrogen starvation media back into nitrogen starvation media (Supplemental Figure 5), suggesting that the change in nutrient abundance, and the associated carbonosome depolymerization, facilitated the luminescence burst observed in WT cells. This transient elevation of intracellular ATP/NADPH concentrations in WT populations aligned temporally with the burst in translation measured via BONCAT (Figure 4B) and the loss of carbonosome-associated BODIPY signal (Figure 2G), suggesting that carbonosome depolymerization might permit simultaneous increases in protein biosynthesis and in key stores of cellular energy and reducing power. The inability of Δ*phaZ* and Δ*phaC1/Z/C2* cells to effectively increase protein synthesis and ATP/NADPH concentrations during these shifts could contribute to the competitive disadvantages observed for these strains in the direct competition assays between WT and each mutant (Figure 3E).

We and others have previously observed that starved *P. aeruginosa* maintain flagellar motility for prolonged periods of starvation and that flagellar components continue to be newly synthesized during growth arrest [20, 51]. We wondered if carbonosome dynamics could impact the motility of growth arrested *P. aeruginosa,* as has previously been suggested for *V. splendidus* [30]. We inspected the motility of WT and Δ*phaC1/Z/C2* populations before, during and after a transition from nitrogen-starvation to carbon-starvation conditions using time-lapse microscopy (Figure 4D). Populations of both strains were highly motile during nitrogen starvation and most motile cells of both strains exhibited high displacement, but the Δ*phaC1/Z/C2* strain had slightly higher fractions of motile cells swimming slightly faster than WT (Figures 4E, F – timepoint 1). Following the transition to carbon starvation media, the percentage of motile cells in populations of both strains dropped substantially, as did the swimming speeds of remaining motile cells (Figure 4E/F – timepoint 2/3). However, in this condition the WT populations retained more motile cells, which swam faster than the Δ*phaC1/Z/C2* cells at the final timepoint sampled (Figure 4E/F – timepoint 3). These subtle distinctions suggest that carbonosomes could impact flagellar motility both during accumulation under nitrogen starvation and during subsequent depolymerization under carbon starvation.

To further investigate the link between flagellar motility and carbonosomes, we probed the ability of the non-motile *flgE:*:tn mutant to accumulate carbonosomes during nitrogen starvation using flow cytometry. We previously observed that the *flgE*::tn mutant accumulated more carbonosome-associated BODIPY signal than the WT during 72 hours of nitrogen starvation [20] but we wondered whether there would be any differences in the timing or magnitude of subsequent changes. We again observed that populations of *flgE*::tn accumulated carbonosomes more quickly and reached a higher maximum carbonosome content than WT but started losing signal again at the same time and to the same relative extent as the WT during later parts of the time course (Figure 4G). Interestingly, *flgE*::tn populations also exhibited lower variance in their carbonosome content (Figure 4H, Supplemental Figure 5). These data revealed interesting interplay among carbonosomes, motility and nutrient abundance. Non-motile cells accumulated carbonosomes more quickly during nitrogen starvation and WT cells that could both produce and later depolymerize carbonosomes retained greater motility following the transition to carbon starvation conditions.

### Carbonosomes impact *P. aeruginosa* tobramycin sensitivity during nitrogen starvation

Growing evidence from many organisms and growth conditions has suggested that reduced metabolic activity correlates with increased tolerance to multiple classes of antibiotics (reviewed in [55]). Our observations that depolymerization of carbonosomes during a transition from nitrogen to carbon starvation could drive higher levels of protein synthesis, intracellular ATP/NADPH, and flagellar motility suggested changes in metabolic activity, even if cells remained unable to grow and divide throughout the transition time course (Figure 4). During repeated fluctuations between the two starvation conditions, the ability to form and depolymerize carbonosomes confers a fitness advantage (Figure 3), but we wondered whether the associated increases in cellular activities could be a liability in the presence of an antibiotic. Therefore, we asked if carbonosome dynamics impacted the activity and viability of growth-arrested *P. aeruginosa* during exposure to tobramycin, an antibiotic commonly used to treat chronic *P. aeruginosa* infections in pwCF [47].

Following 48 hours of incubation in nitrogen starvation media to allow for carbonosome accumulation, WT and carbonosome mutant cultures were exposed to 10µg/mL of tobramycin and sampled longitudinally. The tobramycin MIC value for PA14 under standard Clinical and Laboratory Sciences Institute growth conditions in rich media is reported to be 1 μg/ml, so the 10 μg/ml we used was expected to cause substantial bacterial death. Indeed, the viable CFU counts of all three strains decreased by several orders of magnitude following exposure to 10 μg/ml tobramycin for 24 hours (Figure 5A). However, WT cultures lost viability more quickly, with significantly fewer CFUs recovered than for either carbonosome mutant strain following 4, 6 and 24 hours of exposure to tobramycin. These strain-specific differences in tobramycin survival could also be seen in outgrowth dynamics in liquid media (Supplemental Figures 6B, C). The data suggest that carbonosomes can confer increased sensitivity to tobramycin under nitrogen starvation conditions. Interestingly, under carbon starvation conditions where carbonosomes cannot form, the Δ*phaC1/Z/C2* strain was instead more susceptible than WT to 10µg/mL tobramycin, although both strains were more tolerant in this condition than under nitrogen starvation (Supplemental Figure 6A).

**Figure 5.**
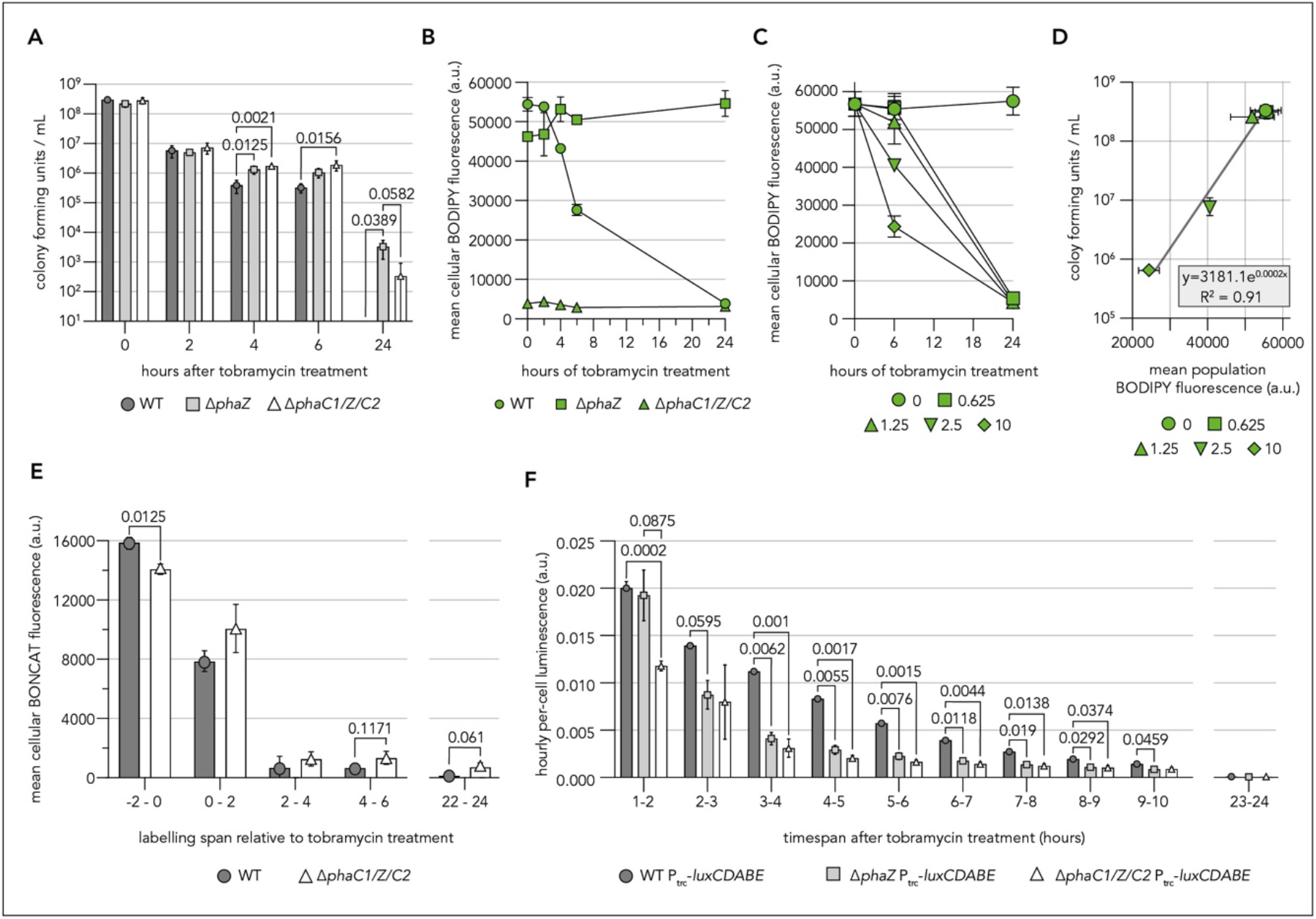
Carbonosomes impact the response of *P. aeruginosa* to tobramycin. Bar graph **(A)** shows CFUs recovered from strains exposed to 10µg/mL tobramycin following 48 hours of incubation in nitrogen starvation minimal media and sampled longitudinally. Line graph **(B)** shows the mean per-cell BODIPY fluorescence in different strains as measured by flow cytometry of populations incubated in nitrogen-starvation minimal media for 48 hours before being exposed to 10 µg/mL tobramycin and sampled longitudinally. Line graph **(C)** shows the mean per-cell BODIPY fluorescence as in B but using only the WT strain and with varying concentrations (µg/mL) of tobramycin. Scatter plot **(D)** shows the correlation between surviving CFUs and the mean cellular BODIPY fluorescence, of populations incubated in nitrogen-starvation minimal media and exposed to varying concentrations of tobramycin (µg/mL) for 6 hours, with trendline shown. Bar graph **(E)** shows the average per-cell fluorescence from BONCAT labelling of cultures incubated in nitrogen starvation conditions for 48 hours before treatment with 10 µg/mL tobramycin. Bar graph **(F)** shows the hourly summed luminescence signal (measured every 6 minutes) from cultures of P_trc_-*luxCDABE* strains incubated in nitrogen-starvation minimal media for 48 hours before treatment with 10µg/mL tobramycin. Luminescence data were normalised to colony-forming unit counts taken immediately prior to application of tobramycin. Means and standard deviations from n=3 biological replicates are shown. One-way ANOVAs with post-hoc Dunnett’s T3 multiple comparison tests were used to assess statistical significance, with only values close to or passing significance thresholds (p < 0.05) being shown.

We next wondered whether tobramycin treatment might be associated with changes to carbonosome content in nitrogen-starved *P. aeruginosa*. We found that exposure to 10 µg/mL tobramycin resulted in the continual loss of carbonosome-associated BODIPY signal over 24 hours in nitrogen-starved WT *P. aeruginosa* (Figure 5B). While further accumulation appeared to halt upon tobramycin exposure, no dramatic loss in signal was observed in Δ*phaZ* populations, suggesting that tobramycin exposure induced active carbonosome depolymerization in WT. By varying tobramycin concentration, we found that the rate of carbonosome depolymerization increased as tobramycin concentration increased, and that complete carbonosome depolymerization was evident following 24 hours of exposure to any tobramycin concentration used (Figure 5C). Furthermore, we found that the remaining carbonosome signal following 6 hours of exposure to tobramycin correlated well with the viable CFUs recovered (Figure 5D). These data suggest that under nitrogen starvation conditions, *P. aeruginosa* responds to tobramycin exposure by depolymerizing material stored in carbonosomes, potentially leading to metabolic activity that is detrimental to survival.

We wondered whether the depolymerization of carbonosomes triggered by tobramycin treatment might be associated with an increase in new protein synthesis or ATP/NADPH pools. Tobramycin targets the 30S subunit of the ribosome, repressing translation to some extent but also causing increased mistranslation of the remaining newly synthesized proteins [56]. If the loss of carbonosome signal during tobramycin treatment is associated with increased synthesis of new proteins, as we observed during the transition from nitrogen starvation to carbon starvation (Figure 3B), this activity could increase mistranslation and reduce tolerance to tobramycin. To investigate this possibility, sequential BONCAT labelling periods were applied to WT and Δ*phaC1/Z/C2* populations before and after treatment with 10 µg/mL tobramycin. Consistent with its mechanism of partial translational repression, tobramycin exposure caused substantial reductions in BONCAT signal, which fell to near-background levels in both strains following 2 hours of exposure (Figure 5E). Small differences were observed between the two strains at some time points but neither exhibited a burst of new protein synthesis concurrent with carbonosome depolymerization. These data suggest that carbonosome depolymerization by WT cells upon tobramycin exposure does not lead to increased mistranslation by substantially stimulating new protein synthesis, and another mechanism must be responsible for the observed decrease in tolerance relative to the carbonosome-deficient mutants.

Because the tobramycin treatment occurred during ongoing nitrogen starvation, which could limit new protein synthesis, we wondered whether carbonosome depolymerization-induced increases in ATP/NADPH levels might instead cause the observed increase in sensitivity to tobramycin. Previous work has suggested that loss-of-function mutations in the NADH dehydrogenase can confer tolerance to aminoglycosides [57, 58]. We reasoned that increased ATP and NADPH pools would be likely to co-occur with increased NADH dehydrogenase activity, which could decrease tolerance.

To examine this possibility, we measured viable CFU counts and luminescence in the luciferase reporter strains following the exposure of nitrogen starved cultures to 10 µg/mL tobramycin. All three strains displayed heightened luminescence (normalized to CFUs) upon tobramycin exposure relative to untreated control cultures (Figure 5F, Supplemental Figure 6). WT and Δ*phaZ* cells quadrupled their luminescence within the first hour analysed, while Δ*phaC1/Z/C2* cells doubled it, suggesting that all three strains quickly sensed and reacted to the addition of tobramycin by increasing their intracellular ATP/NADPH concentrations. However, WT cells maintained significantly higher luminescence than either mutant strain for at least 10 hours following tobramycin exposure. Furthermore, WT cellular luminescence remained above pre-treatment levels for at least 6 hours following tobramycin treatment, while the luminescence of Δ*phaZ* and Δ*phaC1/Z/C2* cells returned to pre-treatment levels within three hours. The increased luminescence of WT cells aligned temporally with the rapid carbonosome depolymerization observed upon tobramycin treatment (Figure 5B/F). Taken together, these data suggest that robust carbonosome dynamics facilitate a sustained increase in cellular ATP/NADPH pools upon exposure to tobramycin.

### Carbonosomes are produced in diverse nutritional contexts and by a variety of *P. aeruginosa* strains

We have employed a reductionist experimental set-up, using defined minimal media, to study carbonosome dynamics and function in growth-arrested *P. aeruginosa*. However, the impact of carbonosome depolymerization on tobramycin tolerance suggests the possibility that carbonosomes could impact antibiotic efficacy in infection contexts. To broaden our investigation of conditions facilitating carbonosome formation, we first investigated an additional nutrient starvation in MOPS minimal media, depriving cells of a source of phosphorus. As with nitrogen starvation, *P. aeruginosa* can survive protracted phosphorus starvation without growing or losing substantial viability (Supplemental Figure 7). While WT and Δ*phaZ* cells quickly accumulated carbonosomes during phosphorus starvation, Δ*phaC1/Z/C2* cells did not (Figure 6A). These data reinforce that carbonosome accumulation occurs when cells have access to excess carbon, but growth is limited by the insufficiency of another essential nutrient.

**Figure 6.**
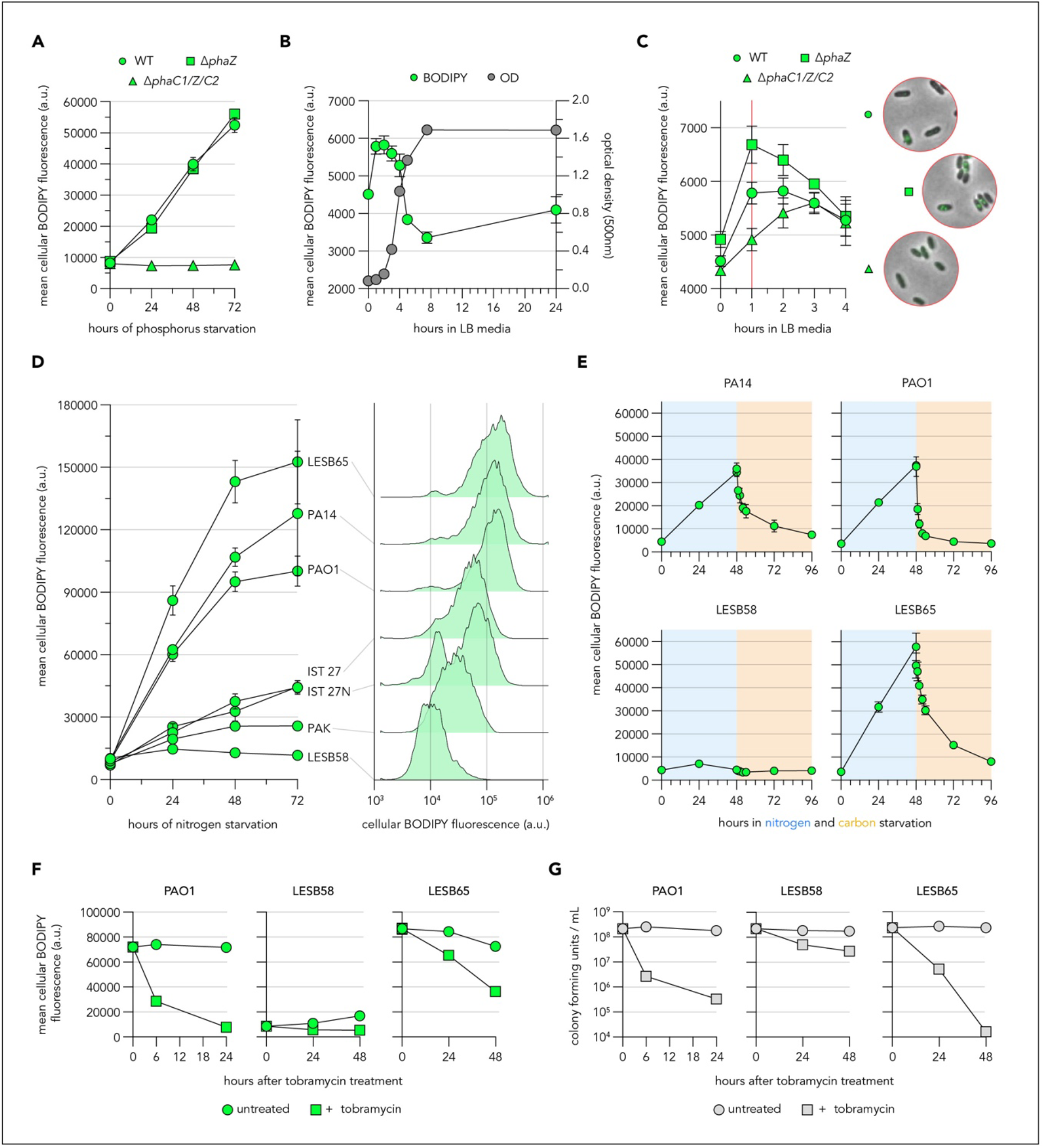
Carbonosomes are produced in diverse nutritional contexts and by a variety of *Pseudomonas aeruginosa* strains. Line graph **(A)** shows the mean cellular fluorescence of BODIPY-stained PA14 cultures incubated in phosphorus starvation minimal media and sampled longitudinally, as measured by flow cytometry. Line graph **(B)** shows both the OD_500_ and the mean per-cell BODIPY fluorescence of WT PA14 cultures grown in LB media and sampled longitudinally. Line graph **(C)** shows the mean per-cell BODIPY fluorescence of PA14 cultures grown in LB media and sampled longitudinally as in B. Representative composite phase-contrast and epifluorescence microscopy images of fixed, BODIPY-stained cells sampled following 1 hour of incubation in LB are also shown. Line graph and histograms **(D)** show the BODIPY fluorescence from cultures incubated in nitrogen-starvation minimal media and sampled longitudinally. Representative per-cell BODIPY fluorescence histograms of cultures starved for 72 hours are shown. Line graphs **(E)** show the mean per-cell BODIPY fluorescence of cultures incubated in and then transitioned from nitrogen-starvation (blue) to carbon-starvation (orange) minimal media and sampled longitudinally. Line graphs show the cytometry-derived mean cellular BODIPY fluorescence **(F)** and the CFUs **(G)** of cultures incubated in nitrogen-starvation minimal media for 48 hours and then treated with tobramycin. PAO1 cultures were treated with 10 µg/mL tobramycin, while LESB58/LESB65 cultures were treated with 40 µg/mL tobramycin. Note the difference in x-axes scales between PAO1 and LESB58/LESB65. Means and standard deviations from n=3 biological replicates are shown.

We next asked if carbonosomes could accumulate in more complex media. We fixed and BODIPY-stained samples from WT cultures over a standard growth curve in LB, from lag phase to stationary phase. Interestingly, we observed a weak spike in average cellular BODIPY fluorescence during the lag phase that dissipated as cells subsequently entered exponential growth (Figure 6B). While some of this signal could be attributed to changes in cellular autofluorescence, the increase in fluorescence during lag phase was more pronounced in Δ*phaZ* populations and less pronounced in Δ*phaC1/Z/C2* populations (Figure 6C) compared to WT. Furthermore, fluorescence microscopy revealed that WT and Δ*phaZ* cells had formed discrete BODIPY-stained puncta within an hour of their resuspension in LB, which more likely reflected small carbonosomes rather than diffuse autofluorescence. Lag phase in nutrient-rich media is a condition where carbon is available in excess and cells are prevented from growing by regulatory or metabolic bottlenecks, although the details of lag phase physiology remain underexplored [59, 60]. The precise role of transient carbonosomes in these circumstances remains unknown, though Δ*phaZ* and Δ*phaC1/Z/C2* mutant strains exhibited no obvious lengthening of lag phase or other growth defect in LB (Supplemental Figure 4). These data suggest that *P. aeruginosa* can produce carbonosomes under a range of non-steady state, slow-or non-growing conditions.

Finally, we examined the ability of *P. aeruginosa* to form carbonosomes in a wider range of genetic backgrounds, including the laboratory strains PAO1 and PAK and a small number of clinical isolates [61, 62]. PAO1 and PAK are representatives of the other major clade of *P. aeruginosa* isolates compared to PA14 [63]. We used nitrogen starvation, which induced stable growth arrest in each strain (Supplemental Figure 7), to promote carbonosome formation and stained cells with BODIPY. Flow cytometry revealed a range of carbonosome content and accumulation rates (Figure 6D), supported by microscopy showing brightly stained puncta (Supplemental Figure 7). The common laboratory *P. aeruginosa* strains PA14 and PAO1 accumulated carbonosomes at similar rates and exhibited comparable population heterogeneity, but PAK showed much lower average levels of carbonosome accumulation and higher heterogeneity. CF clinical isolates IST 27 (a mucoid strain) and IST 27N (a non-mucoid revertant of IST 27) exhibited relatively slow carbonosome accumulation and high population heterogeneity, with clear bimodality in carbonosome content of IST 27N. Interestingly, the CF isolate LESB58 and the closely related LESB65 exhibited dramatically opposing carbonosome phenotypes, with LESB58 populations accumulating no carbonosomes and LESB65 populations achieving the highest carbonosome content of any strain investigated. These data suggest that although most tested *P. aeruginosa* strains could produce carbonosomes, a broad spectrum of phenotypes exists.

We next investigated the ability of PAO1, LESB58 and LESB65 to depolymerise carbonosomes in response to a transition to carbon starvation conditions and to tobramycin exposure. As observed previously, *P. aeruginosa* PA14 depolymerises carbonosomes accumulated in nitrogen starvation media upon the transition to carbon starvation media (Figure 2B, 6E). We found that PAO1 and LESB65 *P. aeruginosa* also responded to transitions with the depolymerization of carbonosomes. Interestingly, PAO1 appeared to depolymerise carbonosomes more quickly, and LESB65 more slowly, than PA14 (Figure 6E, Supplemental Figure 7). LESB58 was unable to produce carbonosomes during nitrogen starvation and so had no carbonosomes to depolymerise in subsequent carbon starvation conditions. We then treated nitrogen-starved cultures of PAO1, LESB58 and LESB65 with tobramycin (doses were adjusted to compensate for differences in tobramycin MICs reported under growth conditions, see legend) and measured CFUs and carbonosome content longitudinally. We observed carbonosome depolymerization in both PAO1 and LESB65 following treatment, while LESB58 produced no carbonosomes (Figure 6F). In PAO1 and LESB65, tobramycin-induced carbonosome depolymerization correlated with dramatic losses in culture CFUs (Figure 6G). Interestingly, LESB58 and LESB65 exhibited substantial differences in tobramycin susceptibility that correlated with their carbonosome content – the carbonosome-containing LESB65 was more susceptible while the carbonosome-lacking LESB58 was less. Although we have not investigated the degree to which the difference in carbonosome content in these strains is directly responsible for the difference in tobramycin tolerance during nitrogen starvation, the trend is in the same direction as that observed for the WT and Δ*phaC1/Z/C2* PA14, where the deletion of genes for carbonosome formation was the only genetic difference.

## Discussion

When subjected to conditions that limit growth, *P. aeruginosa* can store carbon that is available in excess. Here, we have measured benefits and costs of this capacity to store carbon, in the context of fluctuating nutrient limitations and antibiotic treatment. We showed that mutants unable to store carbon in the form of carbonosomes, or to return that carbon to metabolism by depolymerizing carbonosomes, display a strong competitive disadvantage compared to WT when switched repeatedly between nitrogen starvation and carbon starvation conditions. The depolymerization of carbonosomes during shifts between nitrogen and carbon starvation coincided with increases in new protein biosynthesis and ATP/NADPH pools, and WT showed elevated swimming motility following the transition compared to a carbonosome-deficient mutants. These observations suggest that stored carbon can contribute energy and/or biosynthetic substrates to multiple activities that could support survival during ongoing nutrient limitation (Figure 7A). We also observed that this capacity for elevated activity could be a liability during exposure to supra-MIC levels of tobramycin.

**Figure 7.**
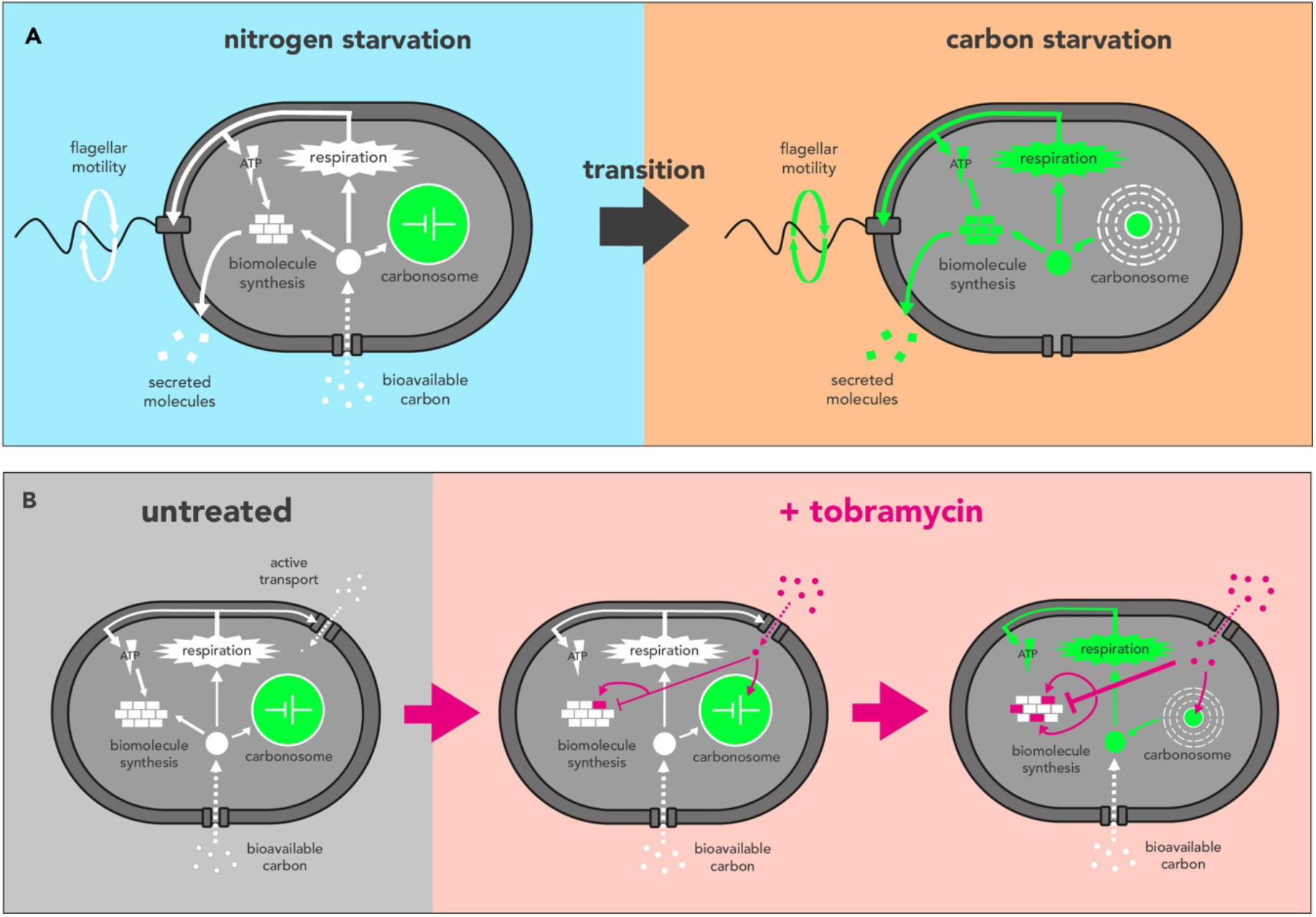
Carbonosomes are intimately wired into the carbon economy of *P. aeruginosa* and can bidirectionally impact fitness in response to stress. Schematics shows the proposed wiring of carbonosomes into physiological activities of *P. aeruginosa.* The flow of carbon into key energetic and biosynthetic pathways is represented by white arrows; The inhibitory effects and impact of tobramycin are represented by red arrows; and the flow of material from carbonosomes into key physiological processes is represented in green arrows. Panel **(A)** shows the accumulation of carbonosomes under nitrogen starvation conditions and the subsequent liberation of carbon to power biosynthesis and respiration upon a transition to carbon-starvation conditions. Panel **(B)** shows the depolymerization of carbonosomes in response to tobramycin, which does not overcome the protein synthesis inhibition imposed by ongoing tobramycin accumulation, but does facilitate increased ATP/NADPH pools.

Unexpectedly, we found that tobramycin treatment during ongoing nitrogen starvation directly triggered carbonosome depolymerization and concurrent increases in ATP/NADPH pools, and that these activities correlated with increased killing by tobramycin (Figure 7B). Finally, we found that cells were able to make carbonosomes in multiple growth-limited states where carbon was in excess, and that the amount of carbonosome accumulated during nitrogen starvation varied greatly across different clinical isolates of *P. aeruginosa*. Together, these results suggest that carbonosomes play important physiological roles during growth arrest, with implications for understanding the biology of growth-arrested bacterial broadly, including in infections.

### Carbonosomes as Dynamic Stores During Growth Arrest

During growth arrest imposed by nutrient limitation, metabolic activity is substantially reduced compared to exponential growth, but new protein synthesis and flagellar motility can be maintained for days or weeks of total starvation. This activity is likely powered by recycling of existing biomolecules, which could appear to represent futile cycles of degradation and re-synthesis from the perspective of the bulk population over time [64, 65]. However, from the perspective of an individual cell in a fluctuating nutrient-limited environment, unpredictable changes in stresses and opportunities might necessitate cycles of breaking down and building biomolecules to repair damage and manage resources available at each moment. Free metabolites can allosterically influence metabolic regulation at multiple levels [66] and can externalize across the cell envelope down a concentration gradient via facilitated diffusion if they accumulate to high levels [67]. Thus, storage polymers that stably sequester excess metabolites are likely to play key roles under non-steady-state metabolic regimes requiring temporally separated cycles of synthesis and degradation. Many questions remain regarding the regulation of production and use of such polymers.

Multiple lines of evidence suggest that carbonosome dynamics are influenced by a complex regulatory and metabolic network that senses cellular status based on multiple inputs. The transcriptional regulator PhaD, encoded adjacent to *phaC1/phaZ/phaC2*, has been shown in *P. putida* to be necessary for WT levels of expression of the *phaC1*, *phaZ*, and *phaC2* genes and for robust carbonosome accumulation [33]. However, a comparison of observed carbonosome dynamics with our previously published nascent proteomics data (Figure 1D) suggest that accumulation and depolymerization of carbonosomes is not controlled only at the level of new expression of the enzymes [20]. Instead, it is likely that activity is also controlled post-translationally, possibly by localization of the enzymes to carbonosomes, interactions with other proteins such as the phasins PhaF and PhaI, or availability of (R)-3 hydroxy fatty acid substrates [68, 69].

We and others have observed dramatic effects on carbonosome accumulation arising from mutations that were not in the carbonosome-associated genes (Figure 1A). Such mutations may contribute to the extreme differences in carbonosome accumulation exhibited by clinical isolates LESB58 and LESB65 (Figure 6D). Previous work identified 137 single nucleotide polymorphisms or small indels between the LESB58 and LESB65 genomes, none of which were in carbonosome-associated genes. Interestingly, LESB65 carries a nonsense mutation in *fleR*, a master regulator of flagellar biosynthesis, and it is non-motile [48]. Our observations of carbonosome synthesis in the *flgE*::tn mutant of PA14 (Figure 4G) suggest that a lack of flagellar motility could contribute to increased carbonosome accumulation. Additional non-synonymous point mutations differentiating LESB65 from LESB58 are found in several other metabolic enzymes and regulators that might influence carbon flux into carbonosomes, such as an acetyl-CoA acetyltransferase, the cytochrome c peroxidase *snr1*, transcription factor *hutC*, and the carbon storage regulator *rsmA* [70–72].

We also previously observed that the global regulator mutants Δ*ptsP* and Δ*ftsH* were completely unable to form carbonosomes, although the mechanisms responsible for this are not known [20, 24]. PtsP is a histidine kinase that is described as a global regulator of carbon/nitrogen balance, with broad impacts on cellular physiology during starvation. Interestingly, its expression has recently been found to be post-transcriptionally regulated by the phasin PhaF, which plays a second role as an RNA-binding protein that regulates translation of a subset of mRNAs in *P. aeruginosa* [37], suggesting the possibility for complex feedback in the regulatory network.

Our results show that carbonosome formation and depolymerization do not strictly depend on specific external carbon sources or nutrient limitations, as we observe formation during nitrogen starvation (Figure 2) or phosphorus starvation (Figure 6A) with succinate as a carbon source, and during lag phase in LB (Figure 6B/C). We also see the net direction of flux shift reproducibly from carbonosome accumulation to depolymerization within the context of ongoing nitrogen starvation (Figure 2G/H), and that depolymerization can be triggered by antibiotic treatment (Figure 5), suggesting sensitivity to a wide range of physiological changes. This is consistent with studies in multiple organisms showing that genetic modulation of many different enzymes in carbon metabolism [31, 73] as well as chemical modulation of electron transport chain fluxes [74] can impact carbonosome accumulation. Together, these observations suggest that PHA dynamics are an output of the entire metabolic network, with opportunities for feedback and crosstalk to buffer metabolism under non-steady-state conditions. These network outputs have been explored to some extent in *P. putida* but further characterization in pathogenic species like *P. aeruginosa* will be an interesting area for future work [75–77].

### Carbonosomes in Infection Contexts

We have not yet explored carbonosome formation in any infection model, but our observations suggest that this may be an interesting avenue for further work. Many studies have suggested that bacteria in infection contexts are unlikely to achieve steady-state exponential growth, instead exhibiting heterogeneous and generally slow growth rates [78–81]. Direct measurements of expectorated sputum from pwCF, who often harbor long-term chronic infections by *P. aeruginosa*, have shown abundant carbon sources, with millimolar levels of glucose, lactate, and amino acids [82, 83]. Growth limitation has instead been proposed to be due to limitations for oxygen, zinc, iron, and/or the stresses of defending against immune cells [84–89], and excess carbon may be available for storage in carbonosomes. It will be interesting to investigate whether laboratory conditions more closely mimicking sputum conditions, or animal infection models, can promote carbonosome formation.

While unexpected, our observation that tobramycin treatment during growth arrest stimulated depolymerization of carbonosomes and a coincident increase in ATP/NADPH levels is also relevant to the CF chronic lung infection context. In pwCF, inhaled tobramycin is frequently used both for control of chronic *P. aeruginosa* infections (maintenance therapy) and for attempts to clear new *P. aeruginosa* infections, especially in young children (antibiotic eradication therapy) [90]. Treatment of chronic infections rarely achieves clearance [47], and antibiotic eradication therapy against new infections also fails 20-40% of the time, but failure is not significantly correlated with tobramycin resistance [91]. This suggests a possible role for phenotypic tolerance of tobramycin in eradication failure. Efforts have been made to identify genotypic and phenotypic correlates of eradication failure in isolates from patients undergoing antibiotic eradication therapy [91, 92], but carbonosome production is not a phenotype that has previously been investigated. Among the small sample of clinical isolates we investigated, we observed large differences in carbonosome formation, and the LESB65 isolate that could form carbonosomes was less tolerant to tobramycin than the closely related LESB58 isolate that could not form carbonosomes. Because carbonosome formation is the output of a complex network with many phenotypic impacts, it is not possible to determine whether carbonosome formation is directly responsible for this difference in tobramycin tolerance. For example, in addition to being non-motile, LESB65 has previously been described to have lower secreted protease and elastase activity [48] and more severe infection phenotypes in a mouse model [50], when compared to LESB58. Other virulence-associated outputs such as secretion of exopolysaccharides could also be affected by carbonosome dynamics [73]. It will be interesting to investigate whether carbonosome formation might be an easily measurable phenotype with predictive power for tobramycin treatment outcomes.

Our exploration of tobramycin sensitivity in isogenic PA14 WT and Δ*phaC1/Z/C2* strains showed that carbonosome formation and depolymerization could directly drive increased sensitivity to tobramycin during nitrogen starvation. This is consistent with previous work investigating mechanisms affecting tobramycin sensitivity. Generally, higher levels of metabolic activity are associated with increased sensitivity to antibiotics [93–95]. However, tobramycin sensitivity has been linked more specifically to increased flux through the NADH-generating reactions of the TCA cycle, and increased cellular respiration, leading to increased PMF (Figure 7B) [96, 97]. Some studies have reported that increased PMF leads to increased tobramycin uptake [98], but others have reported increases in sensitivity that are independent of uptake [58, 97]. The depolymerization of carbonosomes is expected to result in flux of carbon to acetyl-CoA pools, which drive flux through the TCA cycle, leading to increased reducing power in NAD(P)H pools and increased cellular respiration and ATP, as reflected by our luciferase measurements. Our results suggest that growth-arrested bacteria can retain internal metabolic stores in reserve, and that these reserves can be deployed to increase metabolic fluxes and increase antibiotic sensitivity. The mechanisms leading to this deployment are not known and could be an interesting area for future work. Ultimately, improving our understanding of how growth arrested bacteria regulate their stores of metabolites could yield general insights into potentiating antibiotic activity against these cells.

## Data availability

All numeric values used to generate plots are available in Supplemental Data File 1. Raw flow cytometry and microscopy data have been deposited in the European Bioinformatics Institute Biostudies and Bioimages archives respectively. Reviewer links for accessing the raw data are: https://www.ebi.ac.uk/biostudies/studies/S-BSST3423?key=fdeafc3e-9646-459a-bbc9-978c60df69b2 https://www.ebi.ac.uk/biostudies/bioimages/studies/S-BIAD3640?key=b8626aa7-80b6-407c-8395-3c3dfb1ae5d7

## Materials & Methods

### Strain List

| Strain/Organism | Strain ID | Strain Name | Genotype/Description | Origin |
| --- | --- | --- | --- | --- |
| *Pseudomonas aeruginosa*<br>UCBPP-PA14 | MB001 | PA14 WT | WT strain | Gift from Dianne Newman lab [99] |
| | MB058 | PA14 $\Delta phaZ$ | Unmarked deletion of *phaZ* (PA14_66830) | This study |
| | MB059 | PA14 $\Delta phaC1/Z/C2$ | Unmarked deletion of *phaC1-phaZ-phaC2* (PA14_66820, PA14_66830, PA14_66840) | This study |
|  | MB021 | PA14 P<sub>rpsG</sub>-sfGFP | sfGFP under control of P<sub>rpsG</sub> introduced chromosomally at Tn7 site in the background of WT PA14 | Babin et al. 2016 [44] |
|  | MB096 | PA14 P<sub>rpsG</sub>-mScarlet | mScarlet under control of P<sub>rpsG</sub> introduced chromosomally at Tn7 site in the background of WT PA14 | Munro et al. 2026 [20] |
| | MB060 | PA14 $\Delta phaZ$ P<sub>rpsG</sub>-mScarlet | mScarlet under control of P<sub>rpsG</sub> introduced chromosomally at Tn7 site in background of unmarked deletion of *phaZ* | This study |
| | MB061 | PA14 $\Delta phaC1/Z/C2$ P<sub>rpsG</sub>-mScarlet | mScarlet under control of P<sub>rpsG</sub> introduced chromosomally at Tn7 site in background of unmarked deletion of *phaC1-phaZ-phaC2* | This study |
| | MB062 | PA14<br>$\Delta phaC1/Z/C2$<br>$P_{rpsG}$-sfGFP | sfGFP under control of $P_{rpsG}$ introduced chromosomally at Tn7 site in background of unmarked deletion of *phaC1-phaZ-phaC2* | This study |
|  | NA | PA14 *flgE::tn* | *MAR2xT7* insertion within *flgE* (PA14_57960) | Ausubel lab ordered transposon library[100] |
| | MB063 | PA14 $P_{trc}$-*luxCDABE* | *luxCDABE* under control of $P_{trc}$ introduced chromosomally at Tn7 site in WT PA14 background | This study |
| | MB064 | PA14 $\Delta phaZ$<br>$P_{trc}$-*luxCDABE* | *luxCDABE* under control of $P_{trc}$ introduced chromosomally at Tn7 site background of unmarked deletion of *phaZ* | This study |
| | MB065 | PA14<br>$\Delta phaC1/Z/C2$<br>$P_{trc}$-*luxCDABE* | *luxCDABE* under control of $P_{trc}$ introduced chromosomally at Tn7 site in background of unmarked deletion of *phaC1-phaZ-phaC2* | This study |
| *Pseudomonas aeruginosa* PAO1 | MB066 | PAO1 | *P. aeruginosa* strain used in experiments testing a panel of strains for carbonosome accumulation | Gift from Sarah Coulthurst lab [61] |
| *Pseudomonas aeruginosa* PAK | MB067 | PAK | *P. aeruginosa* strain used in experiments testing a panel of strains for carbonosome accumulation | Gift from Daniel Neill lab [49] |
| *Pseudomonas aeruginosa* LESB58 | MB068 | LESB58 | *P. aeruginosa* strain used in experiments testing a panel of strains for carbonosome accumulation | Gift from Daniel Neill lab [49] |
| *Pseudomonas aeruginosa* LESB65 | MB069 | LESB65 | *P. aeruginosa* strain used in experiments testing a panel of strains for carbonosome accumulation | Gift from Daniel Neill lab [50] |
| *Pseudomonas aeruginosa* IST-27 (mucoid) | MB070 | IST 27 | *P. aeruginosa* strain used in experiments testing a panel of strains for carbonosome accumulation | Gift from Daniel Neill lab [49] |
| *Pseudomonas aeruginosa* IST-2N (non-mucoid) | MB071 | IST 27N | *P. aeruginosa* strain used in experiments testing a panel of strains for carbonosome accumulation | Gift from Daniel Neill lab [49] |
| *Escherichia coli* DH5α | MB022 | DH5a<br>pUC18T-miniTn7T- $Gm^R$-$P_{rpsG}$-sfGFP | Harbours pUC18T-miniTn7T delivery plasmid for integrating sequence encoding fluorescent protein into Tn7 site | Babin et al. 2016 [44] |
|  | MB095 | DH5a | Harbours pUC18T-miniTn7T delivery plasmid for | Munro et al. 2026 [20] |
|  |  | pUC18T-miniTn7T- Gm<sup>R</sup>-<br>P<sub>rpsG</sub>-*mScarlet* | integrating sequence<br>encoding fluorescent protein<br>into Tn7 site |  |
|  | MB072 | DH5a<br>pUC18T-miniTn7T- Gm<sup>R</sup>-<br>P<sub>trc</sub>-*luxCDABE* | Harbours pUC18T-miniTn7T<br>delivery plasmid for<br>integrating sequence<br>encoding luminescence<br>machinery into Tn7 site | This study |
|  | MB167 | DH5a<br>pUC18T-miniTn7T- Gm<sup>R</sup>-<br>P<sub>trc</sub>-*NLL-MetRS-*<br>*mCherry* | Harbours pUC18T-miniTn7T<br>plasmid used as vector in<br>construction of lux strains | Babin et al.<br>2017 [101] |
|  | MB073 | DH5a<br>pMQ30-*phaZ*<br>del | Harbours pMQ30 suicide<br>vector for unmarked deletion<br>of *phaZ* gene | This study |
|  | MB074 | DH5a<br>pMQ30-<br>*phaC1/Z/C2* del | Harbours pMQ30 suicide<br>vector for unmarked deletion<br>of *phaC1/Z/C2* genes | This study |
| *Escherichia coli*<br>HB101 | MB010 | HB101<br>pRK2013 | Helper strain carrying<br>machinery for conjugation into<br>*P. aeruginosa* | Gift from<br>Dianne<br>Newman lab<br>[102] |
| *Escherichia coli*<br>SM10 | MB011 | SM10<br>pTNS2 | Helper strain carrying site-<br>specific recombinase for<br>integration into Tn7 site | Gift from<br>Dianne<br>Newman lab<br>[102] |

### Primer List

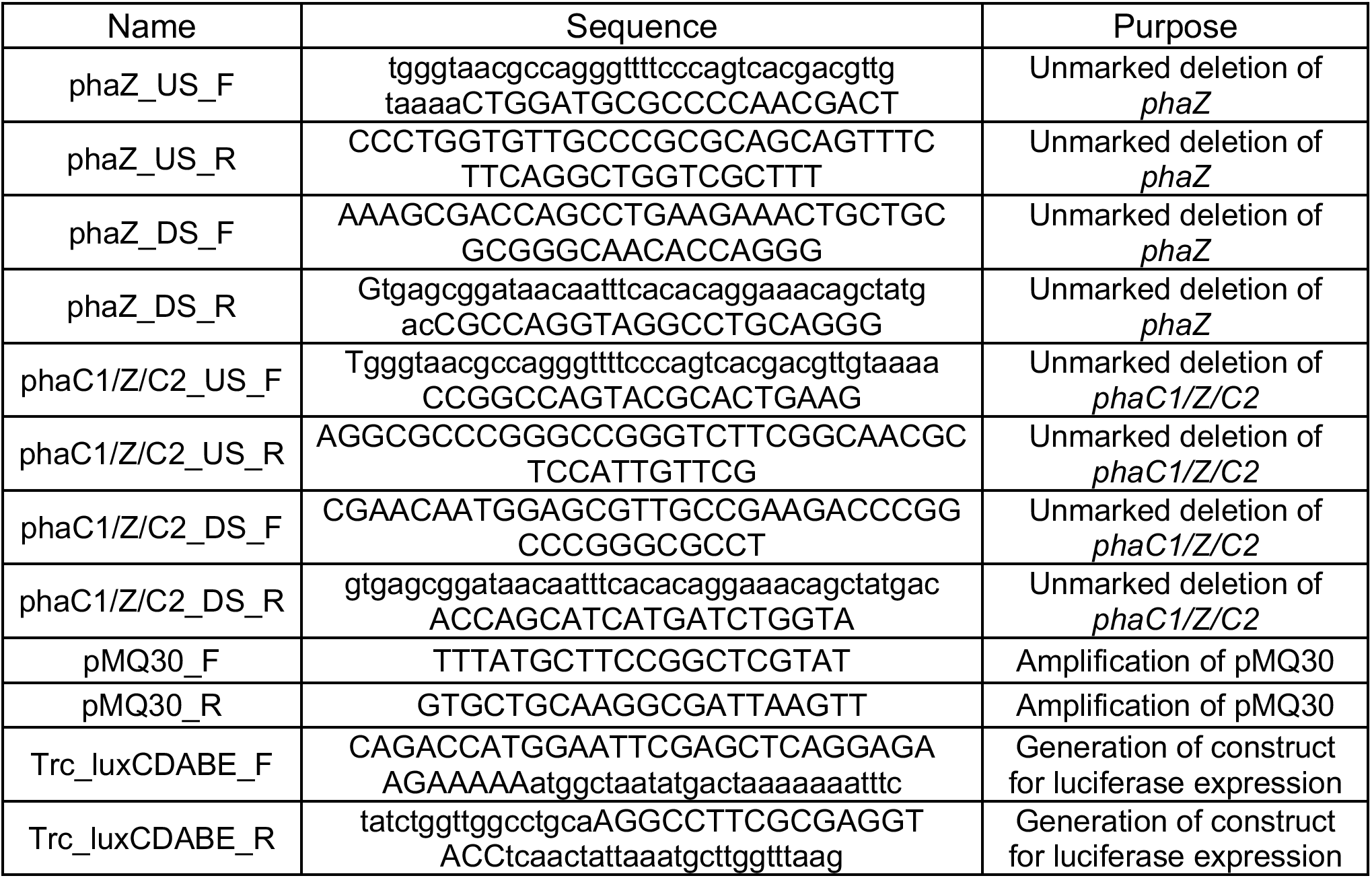
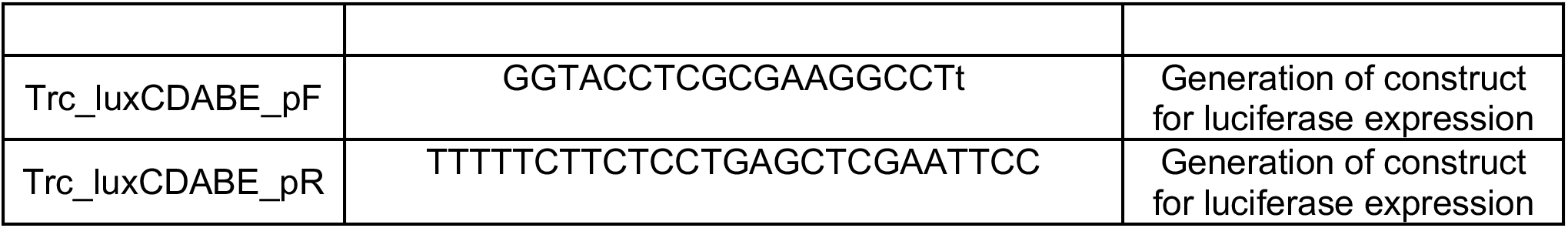

### Strain Construction

Unmarked deletion strains of *phaZ* (Δ*phaZ*) and *phaC1-phaZ-phaC2* (Δ*phaC1*/*Z*/*C2*) were produced by first cloning two 600 bp sequences, encoding sequences upstream and downstream of the relevant genes in either prospective strain, into the pMQ30 suicide vector, using standard methods [103]. Briefly, Gibson assembly was used to combine linearized pMQ30 plasmid (produced by PCR) with the relevant amplified upstream and downstream fragments for each construct, and plasmids isolated from successful DH5a transformants selected on LB agar containing 100 µg/mL gentamycin were confirmed by Sanger sequencing. The pMQ30-*phaZ-*del and pMQ30-*phaC1/Z/C2*-del plasmids were introduced into *P. aeruginosa* UCBPP-PA14 by triparental conjugation using the MB010 as a helper strain. Successful exo-conjugants were selected on VBMM medium containing 100 μg/mL gentamicin as previously described [102] and were then subjected to counter-selection on LB plates lacking NaCl and containing 10% (wt/vol) sucrose. Colonies carrying successful clean deletions were identified by PCR and verified by whole-genome sequencing (SeqCoast and Plasmidsaurus), which also confirmed a lack of spontaneous point mutations. The site of transposon insertion and the absence of consequential point mutations was also validated in the *flgE*::tn strain by whole genome sequencing. The mScarlet and sfGFP reporters under control of the *rpsG* promoter were introduced into the attB site of the relevant clean deletion strains as previously described [20]. To construct strains expressing the *Photorhabdus luminescens luxCDABE* operon under control of the *trc* promoter, the encoding sequence was amplified from plasmid pBEN276 [52] (obtained from addgene) and used to replace the NLL-MetRS encoding sequence within the pUC18-Tn7 P_trc_-*NLL-MetRS*-mCherry plasmid [101] by Gibson assembly. Successfully recovered plasmids were verified by sequencing (Plasmidsaurus) and the luxCDABE-containing cassette was integrated into the Tn7 site of the chromosome of desired PA14 strains using tetra-parental conjugation, as previously described [102].

### Media Conditions, Transitions and Treatments

MOPS minimal media lacking bioavailable carbon, nitrogen and phosphorus sources (50 mM MOPS, 40 mM NaCl, 2.0 mM KCl, 1.0 mM MgSO_4_, 0.1 mM CaCl_2_, 7.5 μM FeCl_2_·4H_2_O, 0.8 μM CoCl_2_·6H_2_O, 0.5 μM MnCl_2_·4H_2_O, 0.5 μM ZnCl_2_, 0.2 μM Na_2_MoO_4_·2H_2_O, 0.1 μM NiCl_2_·6H_2_O, 0.1 μM H_3_BO_3_, 0.01 μM CuSO_4_·5H_2_O) was used as a foundation for all starvation media. Unless otherwise indicated in figure legends, carbon (45 mM sodium succinate), nitrogen (30 mM ammonium chloride) and phosphorus (4 mM KH_2_PO_4_) sources were added as appropriate to create nutrient-replete MOPS minimal media (all three sources) or the desired MOPS starvation minimal media (combinations of two sources). *P. aeruginosa* strains were grown to stationary phase (~24 hours) in lysogeny broth (10 g/L tryptone, 5 g/L yeast extract, 10 g/L NaCl) at 37°C shaking prior to their resuspension in starvation media at OD_500nm_=0.2. Starved cultures were shaken at 37°C in all experiments.

In transitions between starvation media, cells were pelleted (5 mins, 14,000g in benchtop centrifuge) and washed at least once with pre-heated MOPS medium lacking carbon and nitrogen before resuspension in new medium. These transfers can result in the loss of some cells from cultures during transitions (Supplemental Figure 5F), but this is not problematic when measurements are per cell.

Tobramycin working stocks were prepared in culture media and added to cultures following 48 hours of incubation in starvation minimal media; final concentrations are shown in figure legends. In experiments with untreated control samples, an equal volume of tobramycin-lacking media was added to aliquots derived from the same parental culture as treated samples.

### Absorbance Readings and Outgrowth Experiments

The absorbance at 500 nm of cultures and supernatants were recorded using 200 μl aliquots in flat-bottomed clear 96-well plates with a Tecan Spark plate reader. Culture supernatants were clarified by centrifugation (5 mins, 14,000 g in benchtop centrifuge) before aliquoting into the plate. In resuscitation assays (Supplemental Figure 3E), samples (180 μL) from aged cultures were first aliquoted into wells of a flat-bottomed clear 96-well plate. Concentrated solutions of carbon, nitrogen or carbon + nitrogen were then added to wells to give a final volume of 200 μL and a final concentration of 45 mM sodium succinate and/or 30 mM ammonium chloride. In outgrowth assays, strains were diluted from LB overnights (Supplemental Figure 4A/5A) or from aged tobramycin-treated starvation cultures (Supplemental Figure 6B/C) to an OD 500nm of 0.01 in fresh LB. In all resuscitation assays and growth curves, plates were sealed with a BreatheEasy gas-permeable sealing membrane (Diversified Biotech) and placed in a Tecan Spark plate reader pre-warmed to 37°C and set to continuously shake. In all longitudinal experiments, absorbance readings were taken every 6 – 10 minutes.

### CFU Counts

CFUs were assessed by spotting culture samples which had been 10-fold serially diluted in 0.9% NaCl onto LB-agar plates. Plates were incubated at 37°C until colonies became countable, which notably took longer for clinical isolates LESB58 and LESB65.

### Sample Fixation and BODIPY 493/503 Staining

Samples of culture were pelleted (5 mins, 14,000 g in benchtop centrifuge) and washed once with 0.9% NaCl. Samples were then resuspended in 4% paraformaldehyde and incubated for 30 mins at room temperature. Following fixation, samples were washed in 0.9% NaCl and then resuspended in 100 µL 0.9% NaCl and stored in at 4°C until all samples were collected for staining. Samples were pelleted and resuspended in 0.9% NaCl with 1 µg/mL BODIPY 493/503. Samples were incubated in the dark at 37°C for at least 30 minutes prior to use in cytometry or microscopy. For cytometry, samples were diluted in filtered 0.9% NaCl which contained 10 µg/mL DAPI.

### Fluorescence BONCAT Pulse Experiment

Small aliquots (1-2 mL) of culture were amended with 1 mM azidohomoalanine (AHA, Tocris Bioscience). Following labelling, cultures were pelleted (5 mins, 14,000g in benchtop centrifuge) and washed with 0.9% NaCl to remove residual azidohomoalanine. 0.9% NaCl was used to dissolve all subsequent reagents and used to wash cells after each step. Cells were fixed with 4% paraformaldehyde and then permeabilized in 70% ice cold ethanol for 30 minutes. Samples were resuspended with 100 mM iodoacetamide and rotated at 37°C for at least 30 minutes in the dark to alkylate free cysteines. Samples were then pelleted and incubated with 25 μM DBCO-AlexaFluor-488 or DBCO-AlexaFluor-647 (Jena Bioscience) at room temperature for 30 minutes in the dark and were finally washed three times in 0.9% NaCl to remove residual dye. An AHA-free control was used for each condition (pre- and post-perturbation), strain and time point to determine background signal, with the average intensity of this control subtracted from those of actively labelled samples in analyses.

### Flow Cytometry and Analyses

Fixed, stained samples were analysed on a Novocyte (Agilent) flow cytometer. Pulse height measurements were used for all fluorescence measurements at the following wavelengths: BODIPY and AlexaFluor-488 fluorescence 488 nm ex, 530/30 nm em; DAPI 405 nm ex, 445/45 nm em; AlexaFluor-647 640 nm ex, 675/30 nm em. Cytometry data was imported and analyzed in FlowJo V10.8.1 (Becton Dickinson). Populations were first gated based on forward-scatter and side-scatter profiles and then on DAPI signal to identify bacterial cells. Resultant gated populations (4,000-11,000 cells for each biological replicate) were used to produce fluorescence histograms in FlowJo and Adobe Illustrator, and mean and CoV for each population was calculated in FlowJo.

### Microscopy and Image Analyses

Except for motility analyses, all imaging was done using bacteria immobilized on agarose pads. For each agarose pad, a 65 μL gene-frame (ThermoFisher) was filled with 1% agarose and the top surface flattened with a second slide. Sample droplets (0.5 μL) were spotted onto the agarose. All images and videos were acquired using a Nikon Eclipse Ti2 microscope fitted with a 60X phase contrast objective (Nikon-MRD31605); a TIPA slide adaptor (Oko Labs); a Nexus optical breadboard (Thorlabs); a camera (Teledyne photometrics Prime BSI - SN A21b204004) and Spectra light engine (Lumencor - 80 10039). Fluorescence of interest was recorded with the following settings; BODIPY (470 nm LED ex, 510 nm filter em); DAPI (395 nm LED ex, 435 nm filter em); mScarlet (575 nm LED ex, 700 nm filter em) and sfGFP (470 nm LED ex, 510 nm filter em). Consistent phase-contrast settings were used across all experiments, while fluorescence settings were calibrated either to the brightest sample within each experiment for the longitudinal quantification of fluorescence intensities (Supplemental Figure 1) or were calibrated within each individual sample to achieve maximum or clean signal for discerning localization (Figure 2). All microscopy images were processed in ImageJ. Fluorescent channels were aligned to that of the phase using the ImageJ macro Align_Fluor_Channels, written by Norbert Vischer in 2017. The ImageJ plugin MicrobeJ was then used to identify individual cells in the phase contrast channnel and to quantify average pixel intensity per cell for BODIPY staining [104]. MicrobeJ was also used in fluorescence localization experiments to extract cellular BODIPY fluorescence intensity profiles across the defined mid-line of cells. The fluorescence profiles extracted from populations were then averaged and normalized to create representative fluorescence traces (Figure 2F). These localization traces were derived from three distinct replicate cultures, each represented by 58-300 cells. The area occupied by each cell during nitrogen starvation was also quantified using MicrobeJ.

### Transition Competition Assay

Fluorophore-expressing cultures of interest were grown for 24 hours at 37°C shaking in LB media before being washed and resuspended in nitrogen starvation MOPS minimal media at an OD of 0.2. For each competition, strains of interest labelled with opposing fluorophores were then mixed 50:50 and incubated at 37°C shaking. Mixed cultures were incubated for 48 hours in nitrogen starvation MOPS minimal media before being washed and resuspended in carbon starvation MOPS minimal media (considered 1 transition). Following another 48 hours of incubation, cultures were transitioned back to nitrogen starvation MOPS medium, and the cycle was then repeated. Immediately after cultures were first mixed, and at every other subsequent transition, approximately 0.5 μL culture was spotted onto an agarose pad for microscopy as described above. MicrobeJ was used to measure the percentage of cells carrying each fluorophore at each time point. For all competitions, between 110-790 cells were counted to obtain the percentage occupancies in three biological replicate mixtures at each timepoint.

### Cellular Displacement Analyses of Bacterial Motility

Small samples (5-10 μL) were taken from cultures and spotted onto the centre of a 65 µL gene frame (ThermoFisher) attached to a glass slide without agarose. A coverslip was then adhered to the top of the gene frame, squashing the spot. Focusing just below the coverslip, samples were immediately filmed using the phase-contrast channel of a Nikon Eclipse Ti2 microscope. All films were 30 seconds in length and contained 91 frames (3 fps). The ImageJ plugin TrackMate was used to analyse phase-contrast films by identifying individual cells and tracking their displacement between frames of the film [105]. For each film, the data from cells which tracked for less than 50 frames or cells which exhibited extreme displacement indicative of faulty tracking were discarded. Cells were considered motile if they moved more than 1 micron during the 30 second films. Displacement experiments (Figure 4E/F) used the mean percentage motility or cell-speed calculated from six videos derived from three biological replicate cultures, with between 280-2100 cells analysed per video.

### Luminescence Assays

Aliquots (200 µL) of culture were placed in wells of a flat-bottomed opaque 96-well plate. The plate was then sealed with a BreatheEasy gas-permeable sealing membrane (Diversified Biotech) and placed in a Tecan Spark plate reader pre-warmed to 37°C and set to continuously shake and record luminescence every 6 minutes. Values represent 3 biological replicates and were summed over each hour (10 measurements), then divided by the number of CFUs taken before and after a nutrient transition (Figure 4, Supplemental Figure 5). In the case of antibiotic-based experiments, luminescence data was normalised to CFUs taken immediately before the application of the antibiotic (Figure 5).

Although BreatheEasy membranes were vital in preventing the evaporation of samples, we noted that cultures experienced a rapid initial loss of luminescence following their transfer to and sealing within a plate. This response, lasting roughly an hour, was likely reflective of changes to oxygen availability caused by the membrane, and occurred consistently in all strains and replicates. Analyses of luminescence data ignored this initial dynamic response and started one hour after cells were added to the 96-well plate.

### Analyses of Nascent Proteomic Data

Nascent proteomic data was re-analyzed from a previously published dataset [20]. This study used identical starvation media conditions, transitioning methodology and time frames as here. The proteins shown in figure 1 were broadly grouped by enzymatic/structural association with carbonosomes (top) or by contribution to relevant β-oxidation (middle) or *de novo* fatty acid synthesis (bottom) pathways.

### Experimental Design and Statistical Rationale

In the context of frequently employed starvation experiments, biological replicates were defined as parental LB cultures derived from a distinct colony. In bar charts and line graphs, the plotted value is the mean of the biological replicates and error bars represent standard deviation. Flow cytometry histograms depict a single representative replicate. Statistical tests were performed in GraphPad Prism V.10.1.0. Brown-Forsythe/Welch one-way analysis of variance with post-hoc Dunnett’s T3 multiple comparison tests were performed to calculate p-values for pairwise comparisons among multiple strains, while unpaired Welch’s T-tests were performed to calculate p-values for pairwise comparisons between two strains. Comparisons which yielded p-values below the defined significance threshold of 0.05 were always annotated. P-values slightly above this threshold of significance were also sometimes annotated. The trendline in Figure 5D was calculated in Microsoft Excel. All figures and schematics were produced using a combination of Microsoft Excel, Microsoft PowerPoint, GraphPad Prism V.10.1.0, FlowJo V10.8.1 and Adobe Illustrator.

## Acknowledgments

We are grateful for the strains gifted to us by Dianne K Newman, Daniel Neill and Sarah J Coulthurst. We thank Dan Neill, Lisa Racki, Hanna Eriksson, and Claudia Hemsley for helpful feedback. We appreciate assistance from the University of Dundee Flow Cytometry and Cell Sorting Facility, and Imaging Facility. Funding for this work was provided by the Wellcome Trust (Institutional Strategic Support Fund support to MB [204816/Z/16/Z], Wellcome Trust Studentship funding [218520/Z/19/Z] to FDM); the UK Academy of Medical Sciences (Springboard Award [SBF005/1096] to MB); and the UK Research and Innovation Medical Research Council (Future Leaders Fellowship [MR/T041811/1] and [MR/Z000378/1] to MB).

