## Supplemental Figures for "Polyhydroxyalkanoate carbonosomes impact the activity and viability of growth-arrested *Pseudomonas aeruginosa* during nutrient fluctuations and tobramycin exposure"

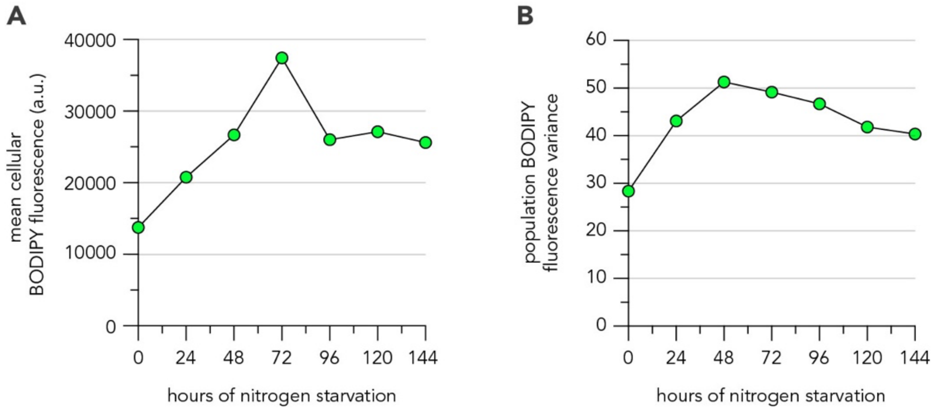


**Supplemental Figure 1 – Quantification of microscopy data**

For quantitative comparison to per-cell fluorescence from flow cytometry, multiple fields per time point shown in Figure 2B were analysed using the ImageJ plugin, microbeJ (see materials and methods for details.) Line graphs show mean cellular BODIPY fluorescence **(A)** and BODIPY fluorescence variance **(B)** extracted from microscopy images, representative images shown in Figure 2B. For each graph, data was extracted from a single biological replicate, with between 430-840 cells analysed at each timepoint.

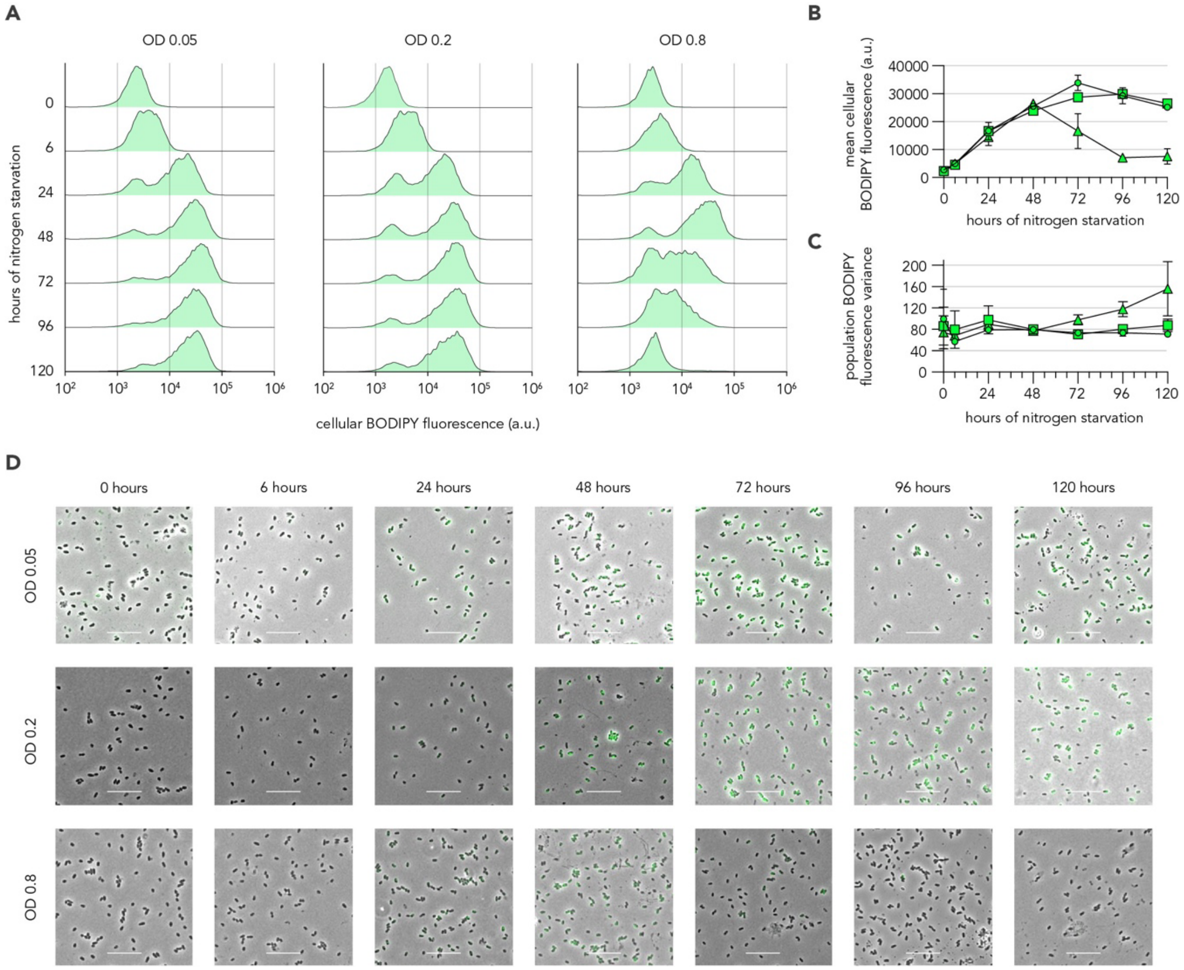


**Supplemental Figure 2 – Impact of culture density on *P. aeruginosa* carbonosome dynamics in nitrogen starvation minimal media**

Histograms **(A)** show the BODIPY fluorescence intensity of populations incubated in nitrogen starvation minimal media at three initial optical densities and sampled longitudinally. Line graphs show the mean cellular fluorescence **(B)** and fluorescence variance **(C)** from fluorescence histograms shown in A. Circles: OD=0.05, squares: OD=0.2, and triangles: OD=0.8. Composite phase-contrast and fluorescence-microscopy images **(D)** show fixed, BODIPY-stained cells from cultures incubated in nitrogen starvation minimal media at defined initial densities and sampled longitudinally. Scale bar represents 10 µm. Each image is from a single replicate deemed representative by cytometry data.

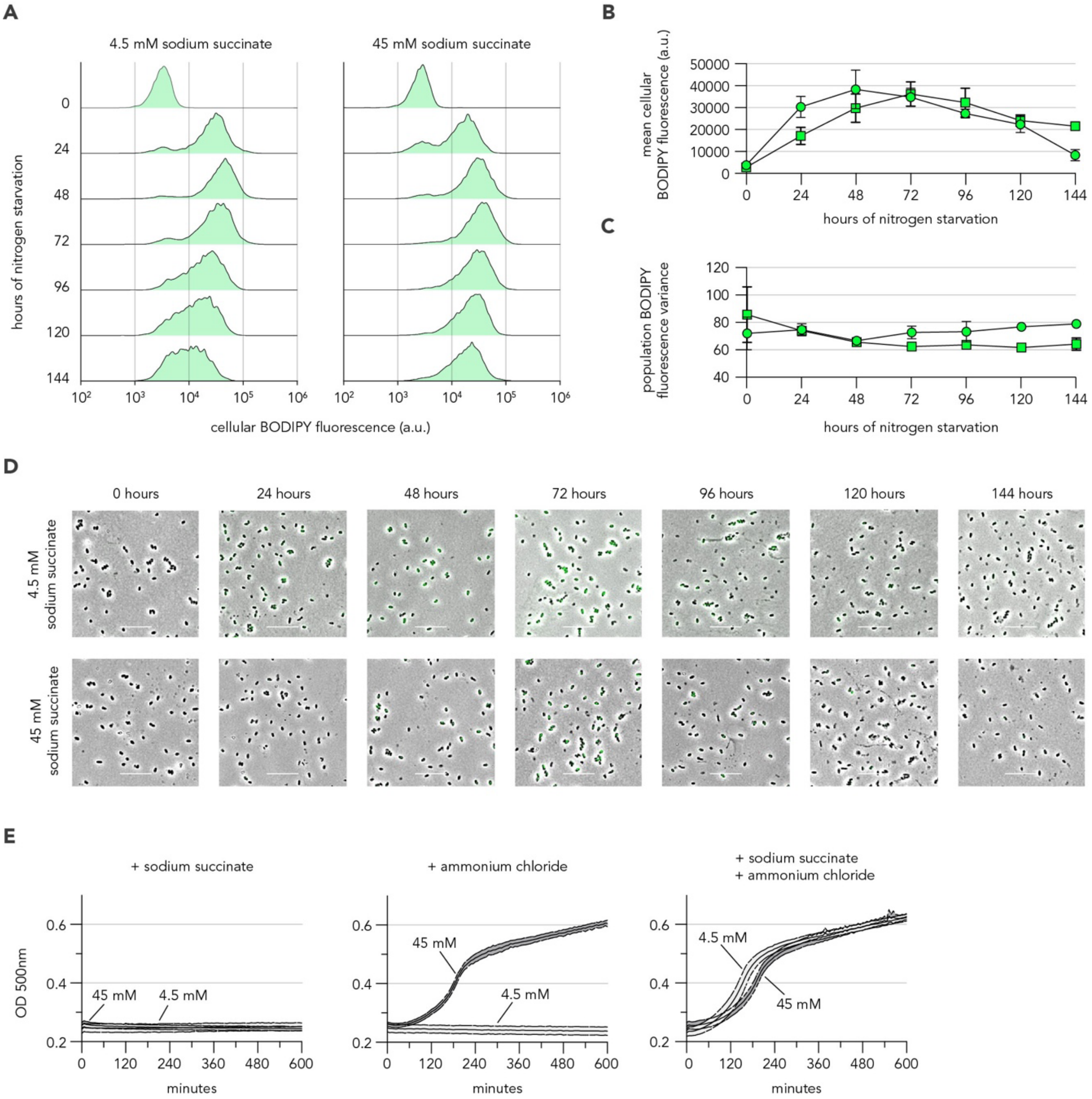


**Supplemental Figure 3 – Impact of carbon-source concentration on *P. aeruginosa* carbonosome dynamics in nitrogen starvation minimal media**

Histograms **(A)** show the BODIPY fluorescence intensity of cells from cultures incubated in nitrogen starvation minimal media with two distinct starting concentrations of carbon source (4.5 mM or 45 mM sodium succinate) and sampled longitudinally. Line graphs show the mean cellular fluorescence **(B)** and fluorescence variance **(C)** extracted from fluorescence histograms shown in A. Circles represent 4.5 mM succinate and squares represent 45 mM succinate. Composite phase-contrast and fluorescence-microscopy images **(D)** show fixed, BODIPY-stained cells from cultures incubated in nitrogen starvation minimal media with two distinct starting concentrations of carbon source (4.5 mM or 45 mM sodium succinate) and sampled longitudinally. Scale bar represents 10 µm. Each image is from a single replicate deemed representative by cytometry data. Line graphs **(E)** show the absorbance-based response of cultures, following 168-hours incubation in nitrogen starvation minimal media with two distinct starting concentrations of carbon source (sodium succinate), to amendment with a carbon source (45 mM sodium succinate - left), nitrogen source (30 mM ammonium chloride - middle) or both (right).

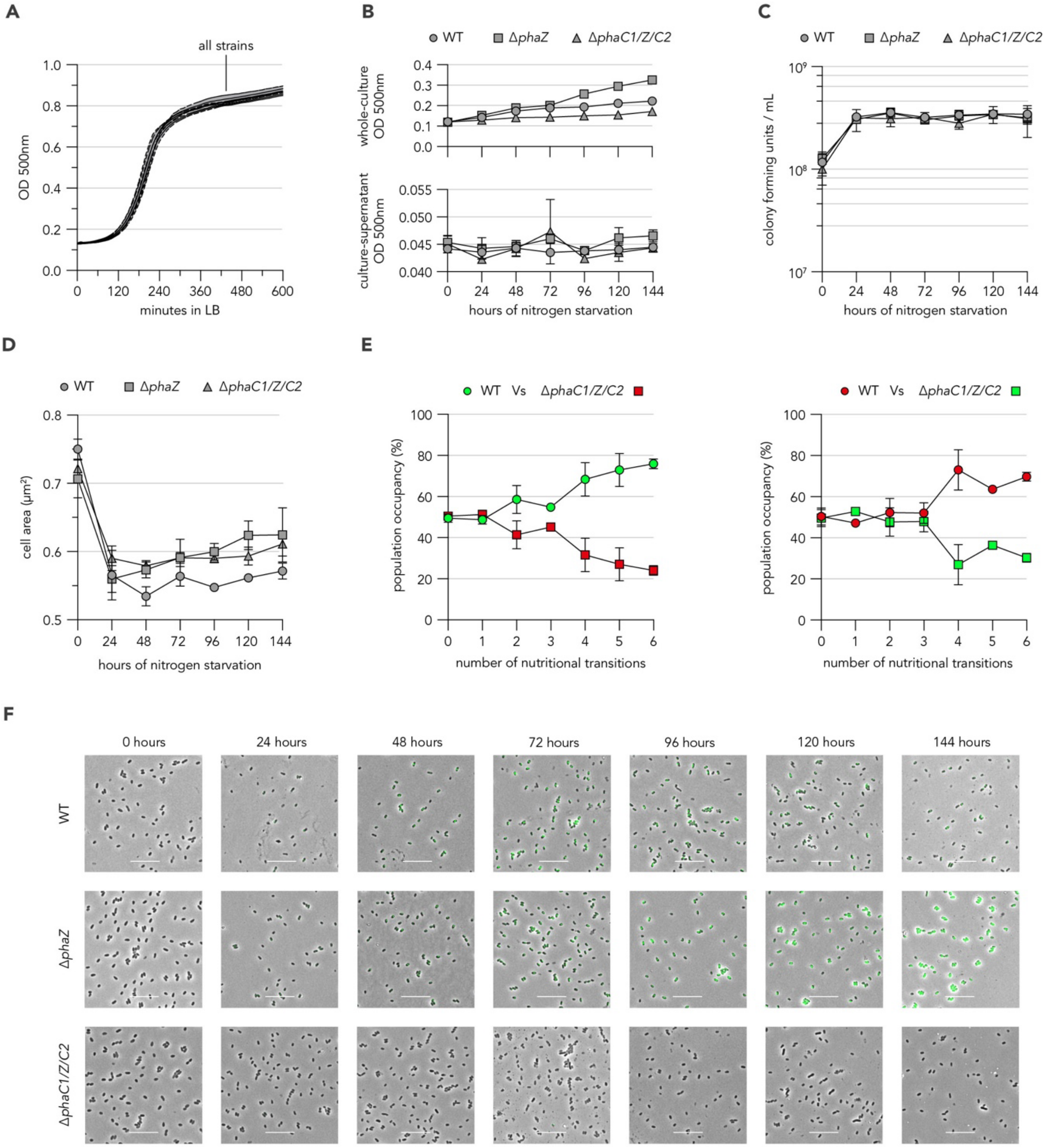


**Supplemental Figure 4 – Auxiliary characterisation of carbonosome mutant strains**

Line graph **(A)** shows optical density growth curves for WT and carbonosome mutant *P. aeruginosa* strains in LB media. Line graphs **(B)** show the whole culture (top) and culture supernatant (bottom) absorbance of WT and carbonosome mutant *P. aeruginosa* strains incubated in nitrogen-starvation minimal media and sampled longitudinally. Note the difference in y-axes scale between graphs. Line graph **(C)** shows the CFUs from WT and carbonosome-mutant *P. aeruginosa* strains incubated in nitrogen starvation minimal media and sampled longitudinally. Line graph **(D)** shows the average cell area, as measured by phase-contrast microscopy, of WT and carbonosome-mutant *P. aeruginosa* cells from cultures incubated in nitrogen-starvation minimal media and sampled/fixed longitudinally. Data is the mean of the average cell area calculated from three distinct replicate cultures, each having between 150-650 cells analysed per timepoint. Line graphs **(E)** show two competitive nutritional transition assays using WT and Δ*phaC1/Z/C2 P. aeruginosa* expressing combinations of distinct fluorescent proteins. Data represent the average percentage of the population occupied by either strain across six transitions between nitrogen-starvation and carbon-starvation minimal media. For each competition, between 60-600 cells were counted to calculate the percentage occupancies in two distinct replicate mixtures at each timepoint. Composite phase-contrast and fluorescence-microscopy images **(F)** show fixed, BODIPY-stained cells from cultures of WT and carbonosome-mutant *P. aeruginosa* incubated in nitrogen starvation minimal media and sampled longitudinally. Scale bar represents 10 μm. Each image is representative of multiple distinct fields and derives from a single replicate deemed representative by cytometry data (Figure 3C).

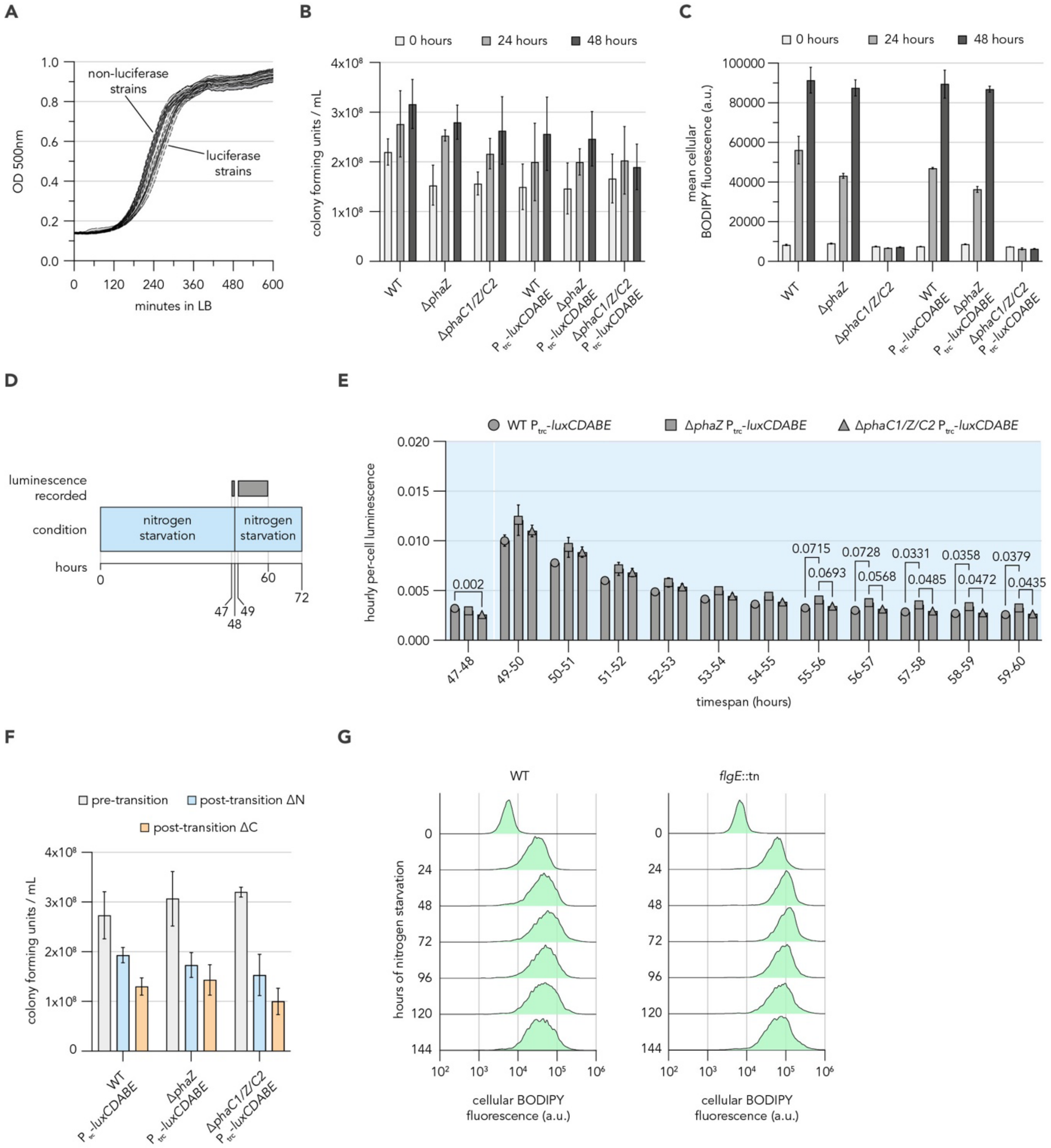


**Supplemental Figure 5 – Auxiliary characterisation of P$_{trc}$-*luxCDABE* and flagellar mutant strains**

Line graph **(A)** shows absorbance-based measure of growth for P$_{trc}$-*luxCDABE* and parental *P. aeruginosa* strains in LB media. Bar graph **(B)** shows the colony forming units retrieved from P$_{trc}$-*luxCDABE* and parental *P. aeruginosa* strains incubated in nitrogen starvation minimal media and sampled longitudinally. Bar graph **(C)** shows the mean cellular fluorescence extracted from cytometry-derived BODIPY-fluorescence histograms of P$_{trc}$-*luxCDABE* and parental *P. aeruginosa* strains incubated in nitrogen-starvation minimal media and sampled longitudinally. Schematic **(D)** shows the timeframe during which luminescence readings were used to interrogate how carbonosome dynamics impacted ATP levels during a transition of P$_{trc}$-*luxCDABE* strains from and into nitrogen-starvation minimal-media. Bar graph **(E)** shows the summed luminescence signal over one hour of measurements made every 6 minutes from cultures of P$_{trc}$-*luxCDABE* strains exposed to the time-course shown in D. Luminescence data was normalised to colony-forming unit counts taken during the incubation of cultures first in nitrogen-starvation minimal media and then following the transition back to nitrogen-starvation minimal media. Bar graph **(F)** shows colony forming units retrieved from P$_{trc}$-*luxCDABE* strains prior to and following transitions from nitrogen-starvation minimal media into new nitrogen-starvation or carbon-starvation minimal media. Histograms **(G)** show the BODIPY fluorescence intensity, retrieved by flow cytometry, of WT and *flgE*::tn populations incubated in nitrogen starvation minimal media and sampled longitudinally.

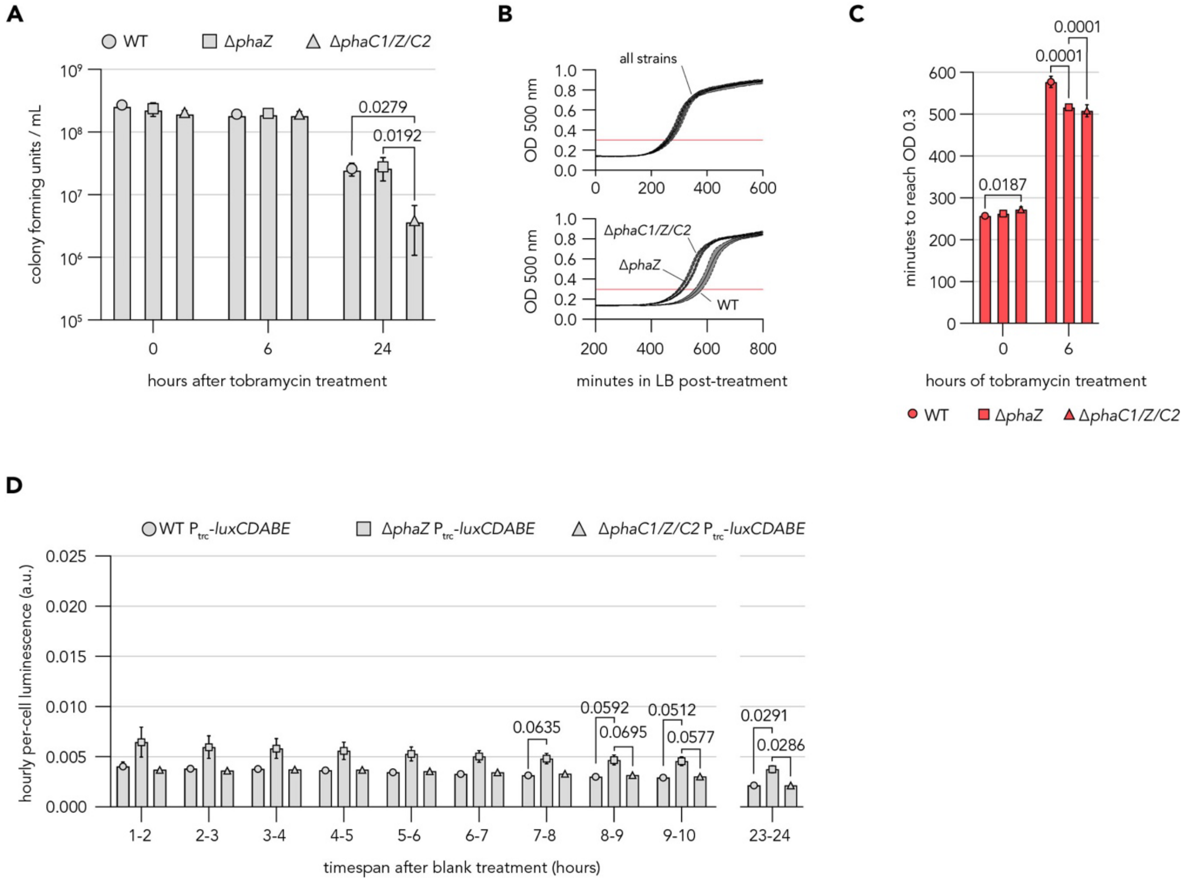


**Supplemental Figure 6 – Auxiliary data from tobramycin susceptibility experiments**

Bar graph **(A)** shows the colony forming units recovered from cultures exposed to 10µg/mL tobramycin during incubation in carbon-starvation minimal media. Line graphs **(B)** show the LB outgrowth of strains incubated for 48 hours in nitrogen starvation minimal media (top) and following an additional 6 hours with 10µg/mL tobramycin (bottom). Note the difference in x-axes scale between each graph. Bar graph **(C)** shows the time taken for LB outgrowths, as shown and treated in B, to reach an optical density of 0.3. Bar graph **(D)** shows the hourly cumulative luminescence signal returned from cultures of $P_{trc}$-*luxCDABE* strains in nitrogen-starvation minimal media upon exposure to buffer lacking tobramycin. Luminescence data was normalised to colony-forming unit counts taken immediately prior to application of buffer.

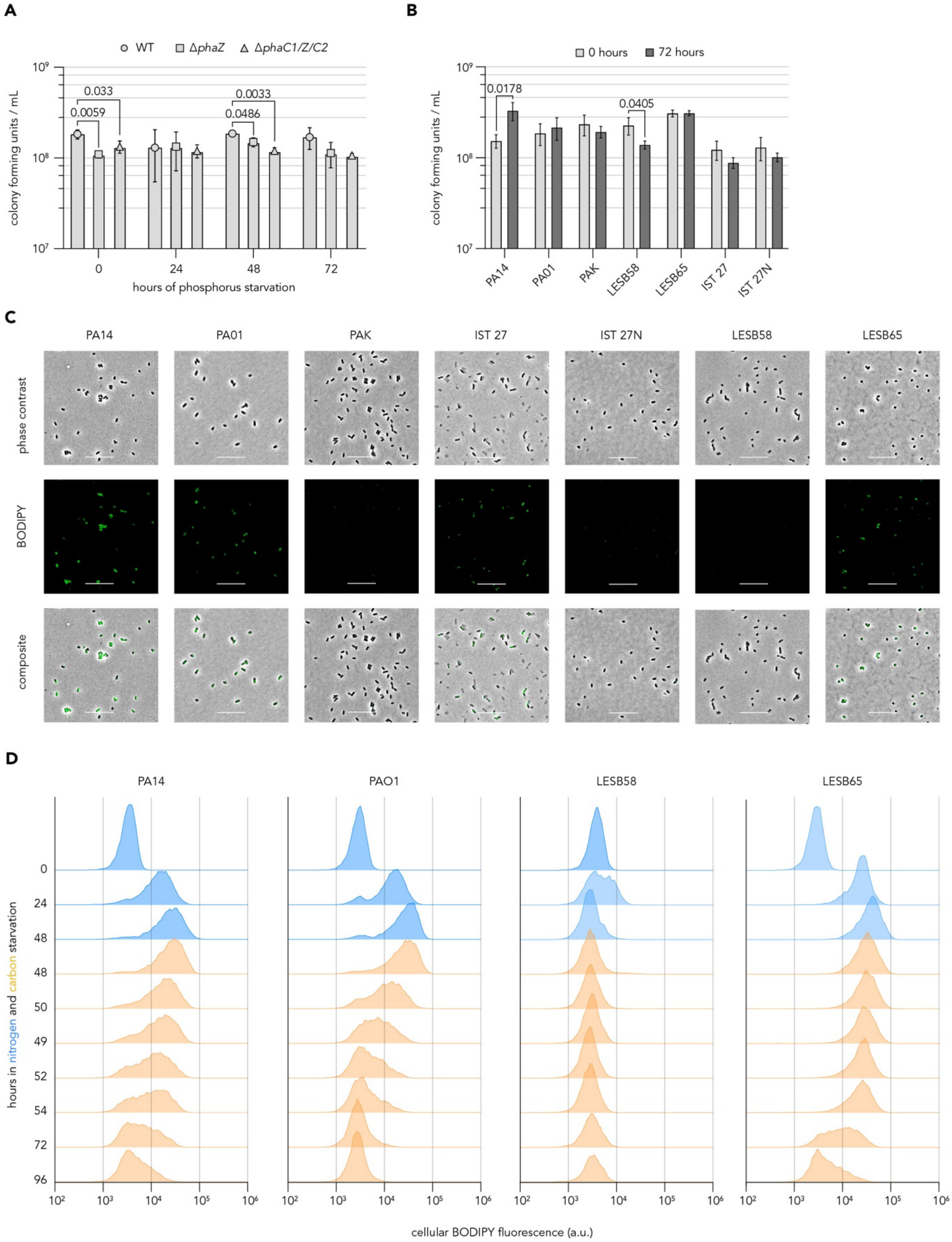


**Supplemental Figure 7 – Auxiliary data from condition and strain investigations**

Bar graph **(A)** shows the colony forming units returned from cultures of WT PA14 and carbonosome-mutant strains incubated in phosphorus-starvation minimal media and sampled longitudinally. Bar graph **(B)** shows the colony forming units returned from cultures of various *P. aeruginosa* strains incubated in nitrogen-starvation minimal media and sampled longitudinally. Phase-contrast and fluorescence-microscopy images **(C)** show fixed, BODIPY-stained cells from cultures of various *P. aeruginosa* strains incubated in nitrogen-starvation minimal media for 48 hours. Scale bar represents 10 µm. Each image is representative of multiple distinct fields and derives from a single replicate deemed representative by cytometry data (Figure 6D). Histograms **(D)** show the BODIPY fluorescence intensity, retrieved by flow cytometry, of populations from various *P. aeruginosa* strains sampled before, during and after a transition from nitrogen-starvation (blue) to carbon-starvation (orange) minimal media.
